# Dynamic and Catalytic Multiphase Coacervates

**DOI:** 10.1101/2025.10.18.683236

**Authors:** Richard Booth, Jiaqi Pei, Jacob Shaffer, Christine D. Keating

**Affiliations:** Department of Chemistry, Pennsylvania State University, University Park, PA 16802

## Abstract

Subcellular organization is critical for myriad cellular functions, including spatiotemporal regulation of biocatalysts. Liquid–liquid phase separation contributes to intracellular organization by providing microenvironments that sequester biomolecules and control their activities. Although catalytic reactions in multiphase coacervate systems are important for living cells, provide functionality to artificial systems, and may aid understanding of how metabolism first arose, they are not well understood. Here we report an artificial multiphase system in which catalytically active droplets containing three distinct coexisting coacervate phases undergo reorganization in response to pH, enabling control over chemical reactions. Catalysis is performed by an inner, polyhistidine-dense coacervate phase with inherent nonenzymatic activity. We show how phase composition, in addition to pH, affects catalytic activity, and how the addition of enzymes that alter local pH provides a means to regulate composition and catalysis. By exploiting competitive interplay between intramolecular self-association and intermolecular complexation among polymers within multiphase coacervate systems, it was possible to (re)organize and tune the reactivity of dynamic synthetic biological systems beyond what is possible for single-phase coacervates. Such approaches could prove valuable in designing artificial cells and related microreactor systems capable of catalyzing and controlling complex chemical and biochemical processes.

## Introduction

Liquid–liquid phase separation (LLPS), and particularly associative LLPS, or coacervation, has recently gained prominence as a paradigm for understanding subcellular organization in biology and for designing non-living compartments with increasingly life-like properties. Coacervate droplets are formed by dynamic, multivalent, interactions whose physicochemical nature can enable responsiveness to external stimuli such as changes in pH,^1,2^ ionic strength,^3,4^ temperature^5,6^ or illumination.^7,8^ The locally high concentrations of coacervate components generate distinct microenvironments that accumulate a wide variety of molecular^9^ and macromolecular^10^ cargoes and can accelerate biochemical reactions.^11–13^ As the number of molecular components is increased, additional coacervate phases are often observed.^14–17^ The wetting properties of coexisting coacervate phases depend on their relative interfacial tensions, which are often ordered by their relative phase separation propensity (e.g., salt stability, relative hydrophobicity) of the polymer combinations.^14–17^ Thus, it is possible by choosing polymer combinations based on the strength of non-covalent interactions and relative hydrophobicity to generate hierarchical multiphase droplet systems.^18^ To date, multiphase systems have been demonstrated that can reversibly form and dissolve in response to enzyme-mediated pH changes,^19^ possess selective RNA duplex thermodynamics,^20^ display additional phase behaviors when integrated with a chemical reaction cycle^21^ and facilitate advanced biomolecular self-sorting into distinct groups when exposed to light or chemical stimuli.^22^ Desirable characteristics may also involve the selective uptake of reactants or the exclusion of products, which, when combined with a crowded hydrophobic interior, are known to accelerate a wide range of chemical reactions.^23^ The potential landscape for tuning properties to enhance chemical transformations expands with the introduction of each new phase in a multiphase system. However, effectively harnessing the increased complexity of non-covalent interactions in multiphase systems to catalyze and drive biochemical reactions remains challenging.

Complex coacervation typically involves two or more oppositely charged polymers that interact by ion pairing.^24^ Additional non-covalent interactions are often involved and can modulate LLPS conditions and coacervate properties including, hydrogen bonding, dipole-dipole, cation-π, π-π and hydrophobic interactions.^25^ Functional groups that can participate in aromatic non-covalent interactions, in addition to charge-based interactions, play a particularly important role in LLPS in biological systems. For example, arginine residues have been shown to be a key driver of LLPS in FUS, LAF1, and Ddx4 proteins.^26^ When arginine is replaced with lysine in these proteins, the phase-separation propensity is reduced, even though the net charge is maintained.^27^ Histidine, another amino acid capable of non-covalent interactions through both charge and aromaticity, has not been studied as extensively as arginine. The near-neutral pKa for histidine’s imidazole group coupled with its aromaticity makes this amino acid an interesting candidate to investigate the interplay between the non-covalent interactions that govern coacervation. In addition, histidine moieties are nearly ubiquitous in enzyme active sites with the histidine-serine-aspartate catalytic triad able to catalyze numerous reactions.^28^ The triad plays a prominent role in enzymes such as oxidoreductases,^29^ acyl transferases,^30^ esterases,^31^ and isomerases.^32^ Enzymes containing histidine-based active sites are prominent biocatalysts with numerous industrial applications such as in the production of biofuels^33^ and the cleaning of environmental contaminants such as plastics,^34^ pesticides^35^ and parabens.^36^ However, incorporating histidine moieties into de novo catalytic systems with comparable catalytic efficiencies to native systems remains elusive.

Incorporating polyhistidine (pHis) into complex coacervates offers the possibility to combine the inherent catalytic activity of histidine moieties with the ability of coacervate droplets to accumulate biomolecules and control local microenvironments potentially leading to de novo soft matter catalytic systems performing with kinetics analogous to native enzymes.^37^ To date, studies of histidine-based coacervates have largely focused on His/Tyr-rich squid beak peptide-inspired sequences, (GHGXY)n, where X is variable,^38^ emphasizing their mechanical properties,^38,39^ drug-delivery potential,^40^ and pH-responsive binding to lipid membranes.^41^ Polyhistidine, which lacks the sequence tunability possible for such peptides, is particularly prone to aggregation and precipitation from solution.^42,43^ The hydrophobicity and potential for strong π-π stacking and cation-π interactions of imidazole moiety, while favorable for mechanical stiffness, can work against reactivity by driving aggregation of pHis peptides when their imidazole groups are predominantly in their catalytically-active, uncharged, form. Solubility can be increased by lowering the solution pH to protonate imidazole side chains, which enhances positive charge and hydrophilicity. However, the pH required to achieve this can fall far below biological levels, especially when pHis interacts with other biomacromolecules.^43^ Moreover, acidification negatively impacts function since only the neutral form of the imidazole moiety is catalytically active. We wondered whether incorporating pHis within droplets comprising one or more coacervate phases might provide sufficient tunability of the microenvironment to prevent its aggregation while maintaining catalytic activity.

Herein, we show how pHis can be used to form stimuli-responsive and catalytically active multiphase coacervate droplets. Surrounding the hydrophobic pHis-rich coacervate droplets with additional coacervate phases altered the molecular composition of the inner histidine-rich phase and led to increased stability against pH-induced aggregation of pHis. The phase organization was reconfigurable through changes in pH that could be induced either by direct pH change or by the activity of incorporated glucose oxidase and urease enzymes. We show how pH-sensitive folding of pHis governs the pH-induced phase reorganization and how this process is impacted by the addition of ATP, which can cation-π and π-π stack with the imidazole moiety, disrupting pHis conformational changes. The degree of cation-π and π-π interactions is also likely pH dependent, further contributing to pH-induced phase behavior and catalytic rate changes. Moreover, the presence of additional coacervate phases enhances esterase-like catalytic activity of the pHis phase by accumulating the reaction product away from the active site. Finally, we show how reaction rates can be controlled by modulating the pH using enzymes. Overall, this work demonstrates that relatively hydrophobic single-phase coacervates can be stabilized against aggregation by additional phases, enhancing their inherent-catalytic activity. This paves the way for creating complex multicompartment coacervates capable of advanced chemical processing and complex biocatalysis.

## Results and Discussion

### Phase behavior of histidine-rich multiphase coacervates

In preparing pHis-based complex coacervates, we sought to balance the greater catalytic activity of the neutral, deprotonated form with the greater solubility of the protonated, cationic form. The pKa for free histidine is 6.0,^44^ local microenvironments can shift the pKa for histidine residues to higher or lower values, for example pKa’s ranging from 5.4 to 7.6 have been reported for different His residues within a single protein.^45,46^ Therefore, we chose an initial pH of ∼6, where the pHis should be partially protonated and carry a net positive charge. We began by mixing pHis (polydisperse, 5-25 kDa), with the negatively-charged polyion polyaspartic acid (pAsp, PDI = 1.00-1.20, ∼11.5 kDa) as a partner to generate complex coacervate droplets. Optical microscopy revealed the formation of small (<5 µm) structures that appeared to be small droplets and gel-like aggregates (Supp. Fig. 1a). We hypothesized that the combination of high charge densities and high total charges of both pHis and pAsp contributed to the formation of aggregates. In an attempt to avoid aggregation, we substituted pAsp with a much smaller anion, adenosine triphosphate (ATP disodium salt, 551.14 Da). Indeed, the resulting droplets appeared larger and more spherical, without obvious aggregation (Supp. Fig. 1b).

We therefore chose ATP as the anion to pHis in constructing our initial multiphase droplets. These pHis/ATP coacervates were combined with additional coacervate phases, starting with pLys/pAsp (pLys, PDI = 1.00-1.20, ∼16 kDa). To prevent aggregation of pHis upon contact with pAsp, multiphase coacervates were prepared from pre-formed single-phase coacervates (pHis/ATP, pLys/pAsp), rather than by mixing the individual polymers together in a single step. This method allowed the buildup of one coacervate phase on top of the other in a stepwise fashion without inducing non-specific aggregation (Figure 1). Mixing pHis/ATP coacervates with pLys/pAsp coacervates immediately resulted in two-phase coacervate droplets with an inner pHis-rich core and an outer pLys-rich shell (Supp. Fig. 2).

**Figure 1.**
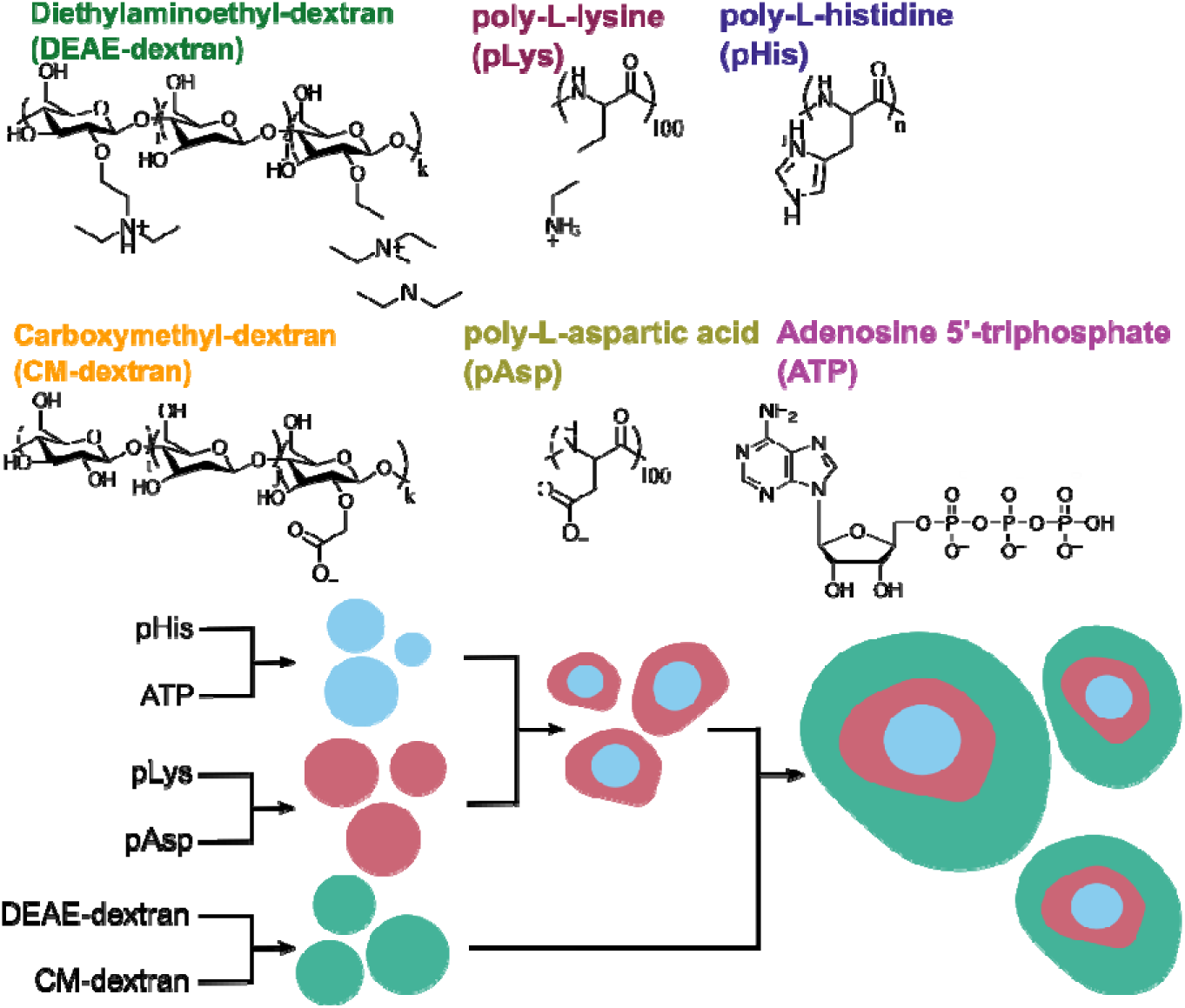
Schematic showing the molecules used in forming the individual single-phase coacervates and how these are combined to form the three-phase multiphase coacervate droplets.

Addition of a third coacervate system, composed of DEAE-dextran (40 kDa) and CM-dextran (polydisperse, 40–60 kDa), generated three-phase droplets. These droplets featured an outer DEAE-dextran–rich phase surrounding a middle pLys-rich phase and an inner pHis-rich core. This organization was revealed by confocal fluorescence microscopy using labeled polycations (Fig. 2a). The final hierarchy consisted of a central pHis–rich core, a surrounding pLys–rich middle phase, and an outer DEAE-dextran-rich layer. Fluorescence intensity profiles confirmed that the three polycations were well segregated with minimal overlap between adjacent phases (Fig. 2b). A separate experiment using labeled polyanions showed that both pAsp and CM-dextran were primarily distributed in the outer and middl coacervate phases, with less pronounced segregation compared to the cationic polymers (Supp. Fig. 3).

**Figure 2.**
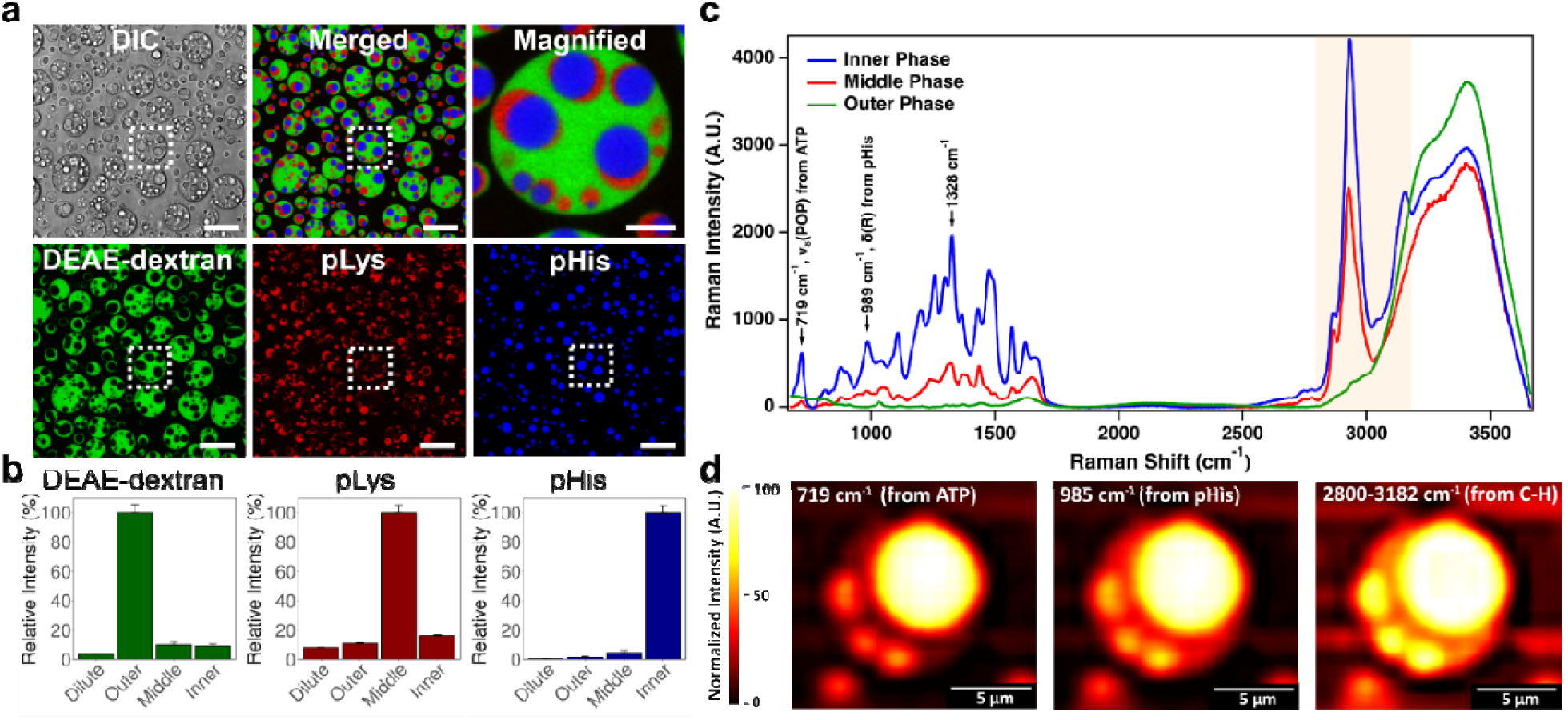
Morphology and molecular distribution in multiphase coacervates. **a)** Confocal fluorescence microscopy images of multiphase coacervates prepared with DEAE-dextran/CM-dextran, pLys/pAsp, and pHis/ATP at pH 5.7. **b)** Comparisons of relative fluorescence intensity for each of the fluorescently labeled cations in the multiphase system (L to R): DEAE-dextran-FITC (green), pLys-Rhodamine (red) and pHis-Alexa Fluor 633 (introduced via a fluorescently tagged 20His-C peptide, blue). Fluorescence intensities are normalized to the maximum signal for each dye, such that the phase with the highest intensity is set to 100%. Intensities in the other phases are expressed as a percentage relative to this maximum. Error bars represent standard deviations of a minimum of 20 droplets. **c)** Raman spectrum of pHis and ATP Raman stretches of multiphase coacervate droplets at pH 6. **d)** MicroRaman maps plotted by characteristic peak: (left panel) ATP distribution represented by phosphate stretching, ν_s_ (P-O-P), at 719 cm^-1^.^47–49^ (middle panel) pHis distribution represented by imidazole ring C-C rocking or scissoring δ (R) at 989 cm^-1^.^50,51^ (right panel) Overall organic content, represented by C-H stretching from 2800 to 3182 cm^-1^. All maps have been normalized based on the area under their corresponding peaks. Scale bars are 10 µm unless otherwise noted.

ATP was not fluorescently labeled due to its small size and lack of suitable functional groups; instead, w used confocal microRaman microscopy to probe its distribution. Both pHis and ATP have characteristic strong Raman-active vibrational modes (Supp. Table 1 and Supp. Fig. 4). microRaman data indicated that ATP was predominantly sequestered in the inner histidine-rich phase, consistent with π-π stacking between the nucleobase and imidazole rings (Fig. 2c, Supp. Fig. 5-6). These measurements also confirmed the location of pHis as mainly in the inner coacervate phase and the same as previously observed by fluorescence intensity data taken via confocal microscopy (Fig. 2a-b). An optical image of a multiphase droplet shows good agreement between the microRaman map and the observed droplet structure (Supp. Fig. 5). microRaman spectroscopy was not able to clearly distinguish between the outer coacervate phase and the surrounding dilute phase, likely due to low concentrations of Raman-active functional groups in these phases.

### pH sensitivity of histidine-rich multiphase coacervates

Since pHis alone is more soluble at low pH, where it is more fully protonated, and tends to aggregate when the pH is above its pKa (pKa ≈ 6-6.5),^45^ its LLPS behavior can be expected to exhibit some pH dependence.^52^ Therefore, we next sought to test the pH sensitivity of the pHis-containing multiphase coacervates. Lowering the pH from ∼5.7 to ∼4.7, thereby increasing the positive charge density on pHis, produced little change in the structural appearance of the multiphase coacervates (DEAE-dextran/CM-dextran, pLys/pAsp, pHis/ATP; Supp. Figs. 7–10), hereafter referred to as the ATP multiphase. Analysi of the relative intensities for labeled cationic polymers showed polyelectrolyte segregation in each phase was also very similar between pH 5.7 and pH 4.7 (Supp. Figs. 7 and 8). We next tested a pH well above the pKa of histidine and found that pHis-rich coacervate droplets were still present in multiphase coacervates (ATP multiphase) even at pH ∼8.5. Some differences were apparent: The pHis-rich droplets were somewhat smaller at the higher pH, with a median diameter of 1.74 µm, 25% smaller in diameter than pH 6.5 (median diameter = 2.35 µm, Supp. Fig. 11). The pHis-rich droplets also sat at the outer edge of the pLys-rich droplets at the higher pH, whereas previously it was contained entirely within the phase, indicating a change in the relative interfacial energies of the phases with different pH.

The relative insensitivity of the multiphase coacervates to differences between pH 5.7 and 4.7 can be rationalized by a coacervate-induced pKa shift of the pHis imidazole groups, in which ion pairing with polyanions stabilizes the protonated (cationic) state to higher pH than observed for free pHis.^53,54^ Although other polyions in this system also have potentially relevant pKa’s, we focus on the pHis protonation state which most directly influences pHis self-aggregation and catalytic activity. Alternatively, it could indicate that the pHis-ATP association is dominated by interactions other than ion pairing that are less affected by changes in pH, such as π–π stacking, or by interactions that can stabilize the protonated imidazole group even at higher pH, such as cation-π interactions. We reasoned that if strong π-π and cation-π interactions between pHis and ATP were responsible for the observed pH insensitivity of the multiphase systems, then substituting ATP with a polyanion unable to engage in π-stacking with pHis imidazoles would increase pH sensitivity. To test this, we replaced ATP with pAsp at the same monomer concentration, while leaving the other components of the multiphase system unchanged; this system is hereafter referred to as the pAsp multiphase. This ATP-free multiphase system indeed showed a strong dependence on pH, with a notable difference in phase morphology and polyion distributions for different pH values. At ∼pH 5.7, the pAsp multiphase system resembled the ATP multiphase system (compare Fig. 3a, b with Fig. 2). However, when the pH of the pAsp multiphase system was lowered to 4.7, the outer DEAE-dextran-rich coacervate phase disappeared, and an additional pLys-rich phase emerged, surrounding the inner pHis-rich phase (Fig. 3c). Fluorescence imaging with labeled polycations revealed similar pLys fluorescence intensities in both lysine-rich phases (outer and middle coacervate phases) at ∼pH 4.7. However, the middle coacervate phase also contained pHis in addition to pLys, whereas the inner phase was predominantly composed of pHis with minimal detectable signal from other cationic components (Fig. 3d). Overall, the ATP-free pAsp multiphase system was similar to the ATP-containing multiphase system at ∼pH 5.7 but different at low pH (∼4.7), losing the outer DEAE-dextran-rich phase and gaining an additional pLys-rich phase (Fig. 3e).

**Figure 3:**
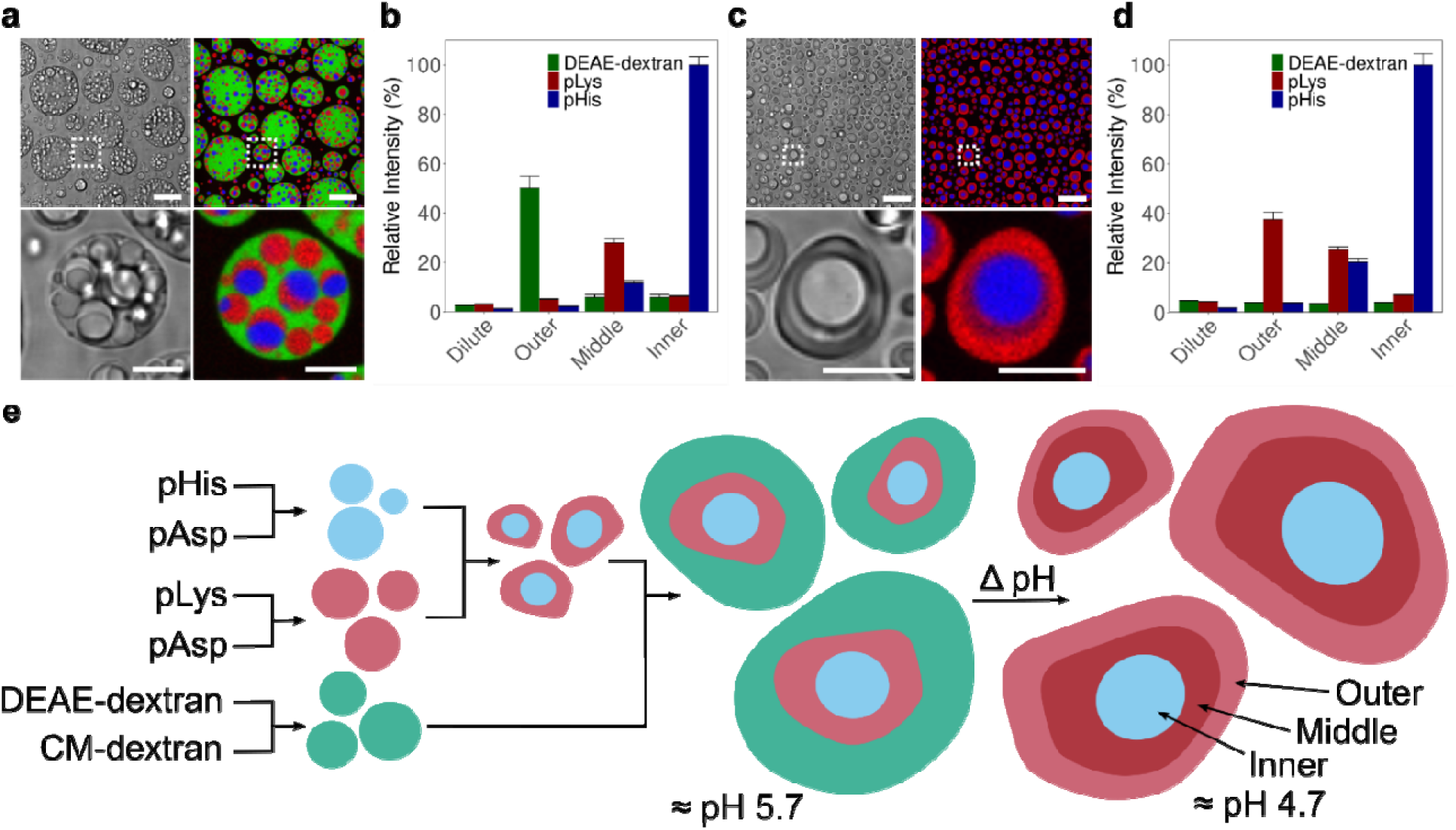
Multiphase coacervate pH sensitivity. **a,c** Confocal fluorescence microscopy images of pAsp multiphase coacervates at ∼pH 5.7 **(a)** and at pH 4.7 **(c)**. **b,d)** Corresponding bar plots of the relative intensity of each of the fluorescently tagged cations DEAE-dextran-FITC (green), pLys-Rhodamine (red) and pHis-Alexa Fluor 633 (blue) in the coacervate multiphase at ∼pH 5.7 **(b)** and ∼pH 4.7 **(d).** Intensities are normalized such that 100% corresponds to the maximum intensity across all dyes and all phases, other dyes and phases are scaled relative to this value. A minimum of 20 measurements were collected per phase. Error bars represent standard deviations. Scale bars are 10 µm (full image) and 2.5 µm for the magnified image that represents the area of the dashed white line box. Error bars in **b, d** represent standard deviations of a minimum of 20 droplets. **e)** Schematic representation of the pH-mediated rearrangement of polyions in the multiphase.

We hypothesized that the pH-responsiveness of the pAsp multiphase system arises from changes in folding patterns of pHis above and below its apparent pKa (≈ 5.0-5.5 in the pAsp system), driven by variations in π-π and cation-π interactions between neighboring imidazole groups as the pH shifts. At ∼pH 4.7, a greater proportion of pHis imidazole groups is protonated. This condition could enable greater electrostatic repulsion between like charges which could promote a more expanded conformation that enhances other heterotypic interactions, such as hydrogen bonding (H-bonding) or electrostatics, between pHis and the other coacervate components (pAsp, pLys, CM-dextran and DEAE-dextran). In contrast, at ∼pH 6, a greater proportion of imidazoles are deprotonated, leading to relatively stronger π-π stacking of neutral imidazoles and cation-π contacts between protonated and neutral imidazole rings, leading to tighter intra- or intermolecular packing and therefore less heterotypic interactions of pHis with the other coacervate components.

To examine this behavior, we recorded circular dichroism (CD) spectra and dynamic light scattering (DLS) measurements for pHis in solution at pH 4.5 and 6.0, either alone or in the presence of each polyanion (Supp. Figs. 12-14). pHis alone undergoes a transition from isolated, random coils to assembled β-sheet-like structures as the pH is increased from 4.5 to 6.0,^55^ consistent with reduced electrostatic repulsion and increased associative interactions due to greater sidechain deprotonation near the pKa (Supp. Fig. 12). Addition of a small amount of ATP or pAsp to pHis (1:100 by monmer) drives self-assembly of pHis; however, the pHis/ATP system is pH-dependent, showing larger assemblies only at higher pH, whereas the pHis/pAsp system forms larger assemblies at both pH values tested (Supp. Fig. 13). The greater propensity for pHis/pAsp association is consistent with the larger size and higher net charge of pAsp compared with ATP.^56^ We argue that ATP more poorly drives pHis assembly at either pH and that the trend in pH dependency observed via DLS reflects that of free pHis. Addition of pAsp to pHis causes only a slight perturbation of the CD spectrum, while ATP addition results in a pronounced reduction of the CD signal at both pH values. This loss of CD signal with ATP reflects disruption of pHis secondary structure (Supp. Fig. 14), likely due to π-π and cation-π interactions between ATP and pHis that interfere with intramolecular imidazole–imidazole contacts stabilizing the secondary structure, even at low pH where ATP is less effective at promoting assembly.^57^ These additional π-π and cation-π interactions, possible with ATP but not pAsp, may underlie the resistance of the ATP-containing multiphase system to strong pH-driven reorganization. From this data, we hypothesized that at low pH, ion pairing and hydrogen bonding dominate, thereby allowing both polyanions to drive LLPS at sufficient concentration. However, at high pH, the increase of pHis π interactions and its reduced net positive charge lowers its affinity for pAsp, leading pAsp to accumulate in other phases thereby driving pH responsiveness. Disruption of pHis π interactions by ATP at higher pH suggests that the loss of ion pairing occurs concurrently with an increase in π interactions, leading to similar distribution of ATP at either pH.

To further explore the role of CM-dextran and pAsp in the pH responsiveness phenomena, we used fluorescently tagged pAsp and CM-dextran to observe their spatial distributions in the multiphase as the pH changes. At ∼pH 4.5, pAsp is predominantly distributed between the middle and outer phases, with higher levels in the middle phase than the outer (Supp. Fig. 15). At this pH, pAsp has ∼2x higher apparent concentration (based on fluorescence intensity) in the middle phase compared to the inner phase and ∼1.5x higher in the outer phase than the inner. The lower relative concentration of pAsp in the inner phase at ∼pH 4.5 likely reflects the dominance of cation-π and hydrogen-bonding interactions between pHis chains and between pHis and pLys at this pH, which reduce the availability of pHis for interaction with pAsp (Fig. 3c,d, Supp. Fig. 15). CM-dextran, on the other hand, is primarily located in the outer phase at ∼pH 4.5, where its concentration is ∼5x greater than in the inner phase (Supp. Fig. 16). This distribution likely reflects its weaker H-bonding capacity with pHis compared to pLys or pAsp, reducing its propensity to form inner-phase complexes under these conditions.

When the multiphase system was prepared at ∼pH 6, both pAsp and CM dextran were largely concentrated in the inner and outer phases, with markedly lower partitioning into the pLys-rich middle phase. The normalized intensity for pAsp in the inner and outer phases is ∼2x higher than in the middle phase, while for CM-dextran, it is ∼5x higher. This observed higher concentration of polyanions in the inner phase runs counter to our hypothesis that ∼pH 6 pHis undergoes strong self-association through cation-π, π-π, and hydrogen bonding interactions, thereby reducing its potential interactions with pAsp and CM dextran. We believe this can be explained through π-π stacking of the aromatic fluorophores covalently bonded to pAsp and CM-dextran and is not necessarily representative of the unlabeled variants of these polymers. We also note that neutral imidizole rings can act as both H-bond donors and acceptors and this could also contribute to higher concentrations of pAsp and CM-dextran in the inner pHis-rich phase ∼pH 6. All together, these findings indicate that the preferred interaction partners of pHis shift substantially depending on pH, with a wide range of heterotypic interactions between pHis polymer chains and also between pHis and the other coacervate constituents (CM-dextran, DEAE-dextran, pLys, pAsp) contributing to the system’s pH responsiveness by forming complexes at some pH values but not others.

### Enzyme-mediated change in pH and morphological changes in multiphase coacervates

To explore the morphological changes in the multiphase system that occur with a change in pH, we used the well-known urease and glucose oxidase (GOx) enzymes to gradually increase or decrease the pH.^19,58^ This transition involves shifting from a system lacking a DEAE-dextran-rich phase at ∼pH 4.7 to a multiphase system with an outer DEAE-dextran-rich phase at ∼pH 6 (Fig. 4a). We first prepared fluorescently labeled GOx and urease to assess their sequestration properties and stability within the multiphase. Confocal microscopy showed that both enzymes were preferentially sequestered in the inner pHis-rich phase, with an intensity approximately 2x higher than in the middle and outer phases (Supp. Fig. 17).

**Figure 4:**
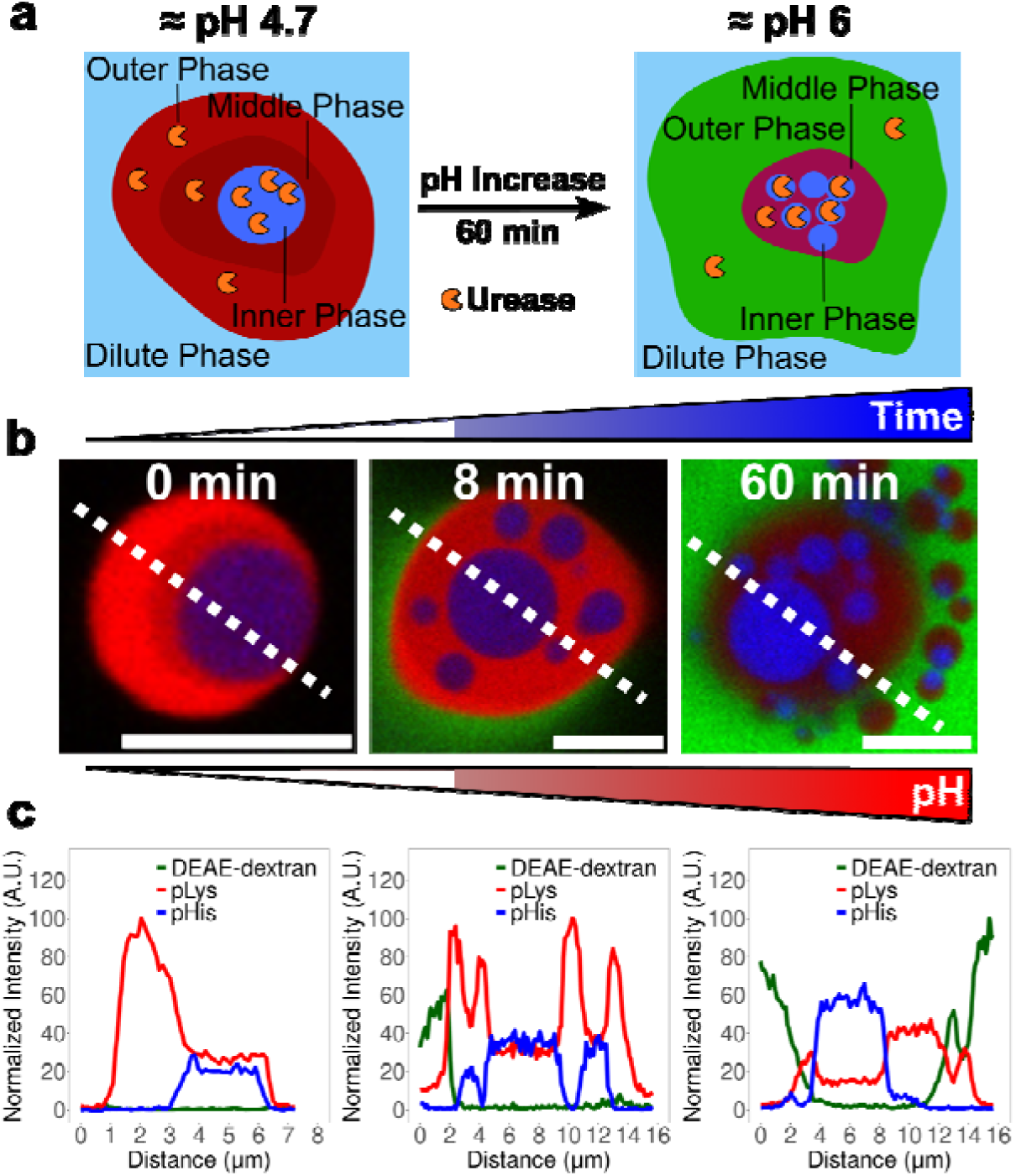
Enzyme mediated phase reorganization in multiphase coacervate droplets. **a)** Schematic showing the morphological differences in the two extremes of pH pAsp multiphase system and the approximate locations of the polyelectrolytes and enzymes. **b)** Confocal microscopy images of the pAsp multiphase over time as the pH increases due to urease mediated production of NH_3_ after the addition of urea (5 mM), dotted line refers to the line profiles. **c)** Line profiles showing the rearrangement of the phases and the change in morphology as the pH increases (plots correspond to dashed lines in **b**). pH of the coacervate mixture was measured via pH probe before and after the reaction to determine the range of the pH change. Scale bars are 10 µm.

Our target pH range using these enzymes spanned the pKa of pHis (≈ pH 5.5), from pH 4.5 to 6 and vice versa (Supp. Fig. 18). The urease reaction was initiated by adding 5 mM urea, and the resulting changes in droplet structure were monitored via confocal microscopy. This revealed that the initial two pLys-rich phases merged into one, followed by the emergence of a DEAE-dextran-rich phase (Fig. 4b,c, Supp. Fig. 19, Supplementary Video 1). As the pH changed, an increase in the number of coacervate droplets was observed, likely due to a change in the hydrophobicity of the histidine-rich polymer, leading to droplet budding (Supp. Fig. 20a, Supplementary Video 1). Droplet settling is not thought to explain this increase in droplets over time because of the duration of the experiment (1 hour) and the high density of the initial droplets which increases the rate at which the droplets settle on the cover slip.

To initiate the reverse reaction and decrease the pH from ∼5.7 to ∼4.5, 15 mM of glucose was added to GOx containing multiphase coacervates. As the pH decreased, the phase rich in DEAE-dextran slowly disassembles and the second lysine-rich phase appears, which is clear from the confocal microscopy images and the line profiles (Supp. Fig. 21a, Supplementary Video 2). It is also noticeable that the mean diameter of the histidine-rich droplets increases due to coalescence as the pH decreases, potentially suggesting a likely decrease in viscosity. This increase in diameter occurs over a period of approximately 5 min, beginning around 20 min after the start of the reaction (Supp. Fig. 20b,c). The 20 min time point in this reaction corresponds to a pH of 5.6 (Supp. Fig. 18), which is around the pKa of histidine.

We further investigated this system by repeating the pH increase reaction using labeled CM-dextran polymer. Initial images at ∼pH 4.5 confirmed CM-dextran was primarily present in the outer pLys-rich phase at this pH (Supp. Fig. 21b, Supplementary Video 3). As the pH increased, the second lysine-rich phase disappeared and was replaced with a CM-dextran-rich phase which comes and goes as the pH increases to ∼pH 6. Given the observed movement of labelled CM-dextran and the general importance of the anions in the rearrangement of the phases as the pH changes, we investigated the movement of both CM-dextran and pAsp as the pH increases. We observed movement of both polyanions during the pH change reaction, which agrees with our previous data (Supp. Figs 15, 16) and suggests that pH-dependent changes in polyanion–pHis interactions play a prominent role in reorganization of the multiphase system (Supplementary Video 4).

### Esterase-like activity in multiphase coacervates

Given the reported inherent catalytic activity of the histidine-rich polymer,^59^ we explored whether our multiphase coacervates could hydrolyze ester-based substrates and act like an artificial enzyme (Fig. 5a). We selected 5(6)-Carboxy-2′,7′-dichlorofluorescein diacetate (CDFDA), which is initially non-fluorescent but converts to the fluorescent product, 5(6)-Carboxy-2′,7′-dichlorofluorescein (CDF) upon hydrolysis of the acetate groups.^60^ We chose this specific substrate due to its high quantum yield at lower pH’s.^61^ First, we tested the sequestration properties of the CDF product by confocal microscopy (Fig. 5b). The fluorescent CDF is predominantly localized in the inner histidine-rich phase, showing a relative intensity ∼2x greater than in the middle and outer phases at both pH ∼4.5 and ∼5.7. Interestingly, the CDF dye also had a ∼2x higher relative intensity in the middle pLys-rich phase at ∼pH 5.7 compared to ∼pH 4.5 (Fig. 5c). Based on the sequestration behavior of the CDF product within the multiphase system, particularly in the pHis-rich phase, we expected the non-fluorescent CDFDA substrate to behave similarly and localize in the catalytically active pHis-rich phase. Upon adding CDFDA at ∼pH 5.7, confocal microscopy confirmed this: the substrate underwent ester-like hydrolysis, initially starting in the inner pHis-rich phase and then diffusing outward into the surrounding phases, with a strong green fluorescence developing in the inner phase over 900 seconds (Fig. 5d,e, Supp. Fig. 22, Supp. Video 5).

**Figure 5:**
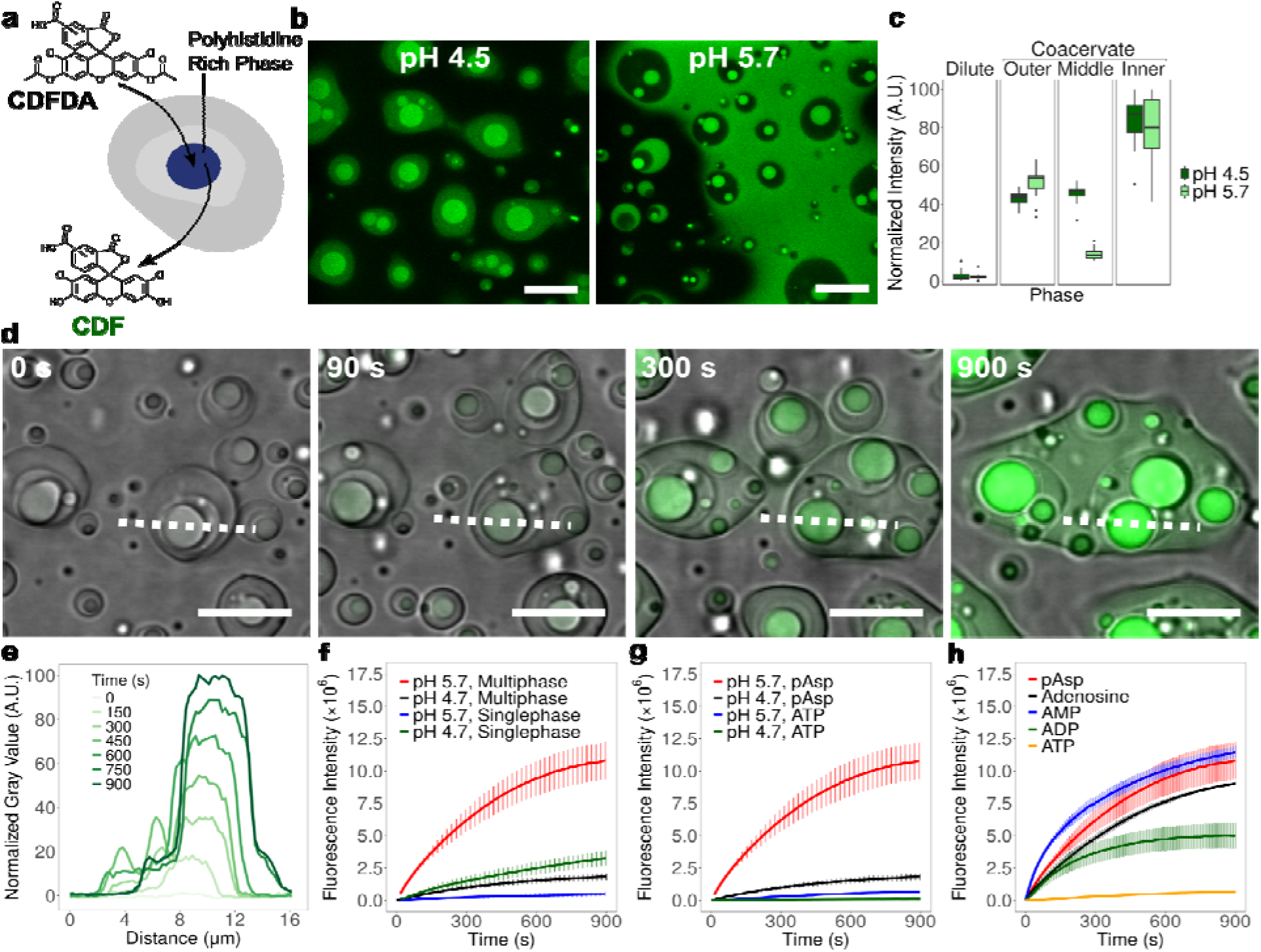
Esterase-like hydrolysis in multiphase coacervates. **a)** Schematic showing the addition of the ester-containing fluorophore (CDFDA) and the subsequent hydrolysis by the inner pHis-rich phase of the multiphase to the fluorescent product (CDF). **b)** Merged confocal microscopy images of the transmitted light (DIC) and green fluorescence channel of CDF sequestration at pH 4.5 and 5.7 for the pAsp multiphase system. The outer phase has coalesced to form a near continuous phase at pH 5.7 **c)** Corresponding box plots showing the normalized intensity (min-max) of the sequestration of CDF in the multiphase system at either pH 4.5 or pH 5.7, each pH was normalized separately and a minimum of 20 intensity values were taken. **d)** Merged confocal microscopy images of the DIC and green fluorescence channel showing formation of the CDF over a period of 900 s in the CM-dextran/DEAE-dextran, pLys/pAsp, pHis/pAsp multiphase system. **e)** Line profile plots showing the increase in fluorescence intensity across the multiphase over the course of the 900 s reaction (plots correspond to dashed lines in **d)**. **f-h** Fluorescence spectroscopy kinetics showing the hydrolysis of CDFDA in the multiphase and single-phase coacervates at pH’s 4.7 and 5.7, the units are arbitrary (λ_ex_ 492 nm, λ_em_ 517 nm) **(f)** the multiphase and single-phase pHis/pAsp coacervates at pH’s 4.7 and 5.7, **(g)** the multiphase in the presenc and absence of ATP at pH’s 4.7 and 5.7 and **(h)** the multiphase in the presence of pAsp, adenosine, AMP, ADP and ATP at pH 5.7. Error bars in **f-h** represent standard deviations of a minimum of 3 under each set of conditions. Scale bars are 10 µm.

In order to get an accurate measure of the kinetics of the reaction by fluorescence spectroscopy, we first tested the pH sensitivity of free CDF product by taking fluorescence spectra of free dye at pH’s 4.7 and 5.7 to calculate a correction factor for pH dependent fluorescence (Supp. Methods, Supp. Fig. 23). Comparison of the rate of hydrolysis in the multiphase at pH 5.7 and 4.7 by fluorescence spectroscopy revealed that the rate of hydrolysis was ∼7x greater at the higher pH (Fig. 5f, Supp. Table 2). This could be because the crowded environment of the multiphase coacervates stabilizes pHis, which is less soluble at high pH but more catalytically active due to the presence of the lone pair on the neutral imidazole ring at these pH levels.

Interestingly, the rate of reaction was considerably faster in the multiphase system than the single-phase pHis/pAsp coacervates, with an apparent average increase in rate of ∼6x across the two pH values (Fig. 5f, Supp. Table 2). A possible explanation for this effect is that the outer phases can sequester the product, effectively removing it from the active site and thereby driving the reaction forward by relieving product inhibition. A further interesting observation is that the hydrolysis rate was faster at the lower pH of 4.7 compared to the higher pH of 5.7 for the single-phase coacervates, which is the opposite trend seen in the multiphase system (Fig. 5f, Supp. Table 2). In addition, control experiments without pHis showed negligible background hydrolysis, even when multiphase coacervate droplets (DEAE-dextran/CM-dextran, pLys/pAsp) were present to enhance the solubility of CDFDA (Supp. Fig. 24 Table 2).

The spectra of free dye at the two pH values also enabled us to calculate how close to completion the coacervate systems are by comparing the fluorescence intensity of the reaction mixture to that of the free dye (Supplementary Table 2). The multiphase system at ∼pH 5.7 reached approximately 20% completion after 900 seconds, which was about six times higher than the ∼3% completion observed at ∼pH 4.7. However, when comparing the multiphase system to free pHis polymer without coacervates, the free polymer reached around 40% completion, approximately twice as fast as the multiphase system (Supplementary Table 2 and Supp. Fig. 25). The lack of rate acceleration in the presence of coacervates could be due to the relatively high substrate concentrations used in all these reactions.^4^ It is also worth noting that pH 4.7 was quicker than pH 5.7 for the free polymer which is opposite to the multiphase system but the same as the single-phase system.

Comparison of the pAsp system and the ATP system showed hydrolysis rates decreased by ∼50x when ATP was present in the system (Fig. 5g). We hypothesized that this was due to ATP’s strong interactions with the pHis imidazole groups through both the adenosine group π-π and cation-π stacking with the imidazole ring and the negative charge of the phosphate electrostatically bonding to the positively charged histidine residues. To further investigate the effect of ATP on the catalytic activity, we compared the effect of ATP with analogues having fewer phosphate groups: ADP, AMP, and adenosine. Fluorescence spectroscopy showed an increase in rate as the phosphates are removed with ATP exhibiting the slowest kinetics and AMP the fastest (Fig. 5h). The AMP system even outperformed the ATP-free pAsp system in the initial timepoints exhibiting kinetics that were ∼2x faster (Fig. 5h). Interestingly, removing all phosphates from ATP did not provide further rate enhancement: adenosine’s product formation curve falls between that of AMP and ADP, indicating other factors in addition to charge play a role in the kinetics (e.g., solubility, impact on phase properties and hydrogen bonding of phosphate oxygens with other components).

To further investigate how pAsp and ATP influence the catalytic activity of pHis, we performed fluorescence recovery after photobleaching (FRAP) on fluorescently labeled pHis in both systems. In the single-phase coacervates FRAP revealed slower recovery in the ATP-containing system compared to the pAsp-containing system, indicating that pHis is more mobile in the presence of pAsp. This increased mobility suggests greater accessibility of catalytically active imidazole groups to interact with the substrate (Supp. Figs. 26, 28a). FRAP for the ATP multiphase and pAsp multiphase coacervates also showed ATP coacervates had slower recovery, potentially indicating that the ATP system is more viscous, and that this increased viscosity could be contributing to the slower kinetics when ATP is present (Supp. Figs. 27, 28b). A smaller difference in the rate of recovery between pAsp and ATP was observed in the multiphase system compared with the single-phase system, indicating that the presence of the multiphase reduced the disparity in recovery rates by enhancing recovery in the ATP system but not in the pAsp system.

### Kinetics of esterase-like catalysis in a changing pH environment driven by urease

Given the strong dependence of reaction rate on pH, we wondered if an enzyme-mediated pH change could act like a switch for the hydrolysis reaction. We incorporated urease to increase pH (Fig. 6a). Indeed, imaging during this reaction revealed an increase in product (CDF) fluorescence as well as the characteristic change in coacervate morphology that accompanies the pH increase (Fig. 6b). As in the enzyme-free reaction (Figure 5), the fluorescent product first appeared in the inner pHis-rich phase before diffusing to the surrounding phases of the multiphase droplets (Fig. 6b,c, Supp. Fig. 29). When the reaction rate was monitored by bulk fluorescence spectroscopy of multiphase droplet suspensions, to our surprise we found that the rate of hydrolysis for the urease-containing system (initial pH = 4.7, increasing due to urease activity) was as high as for the enzyme-free system (pH 5.7), even at early timepoints when the urease-containing system’s pH was still below pH 5 (Fig. 6d). Possible explanations include locally higher pH even at earliest timepoints due to compartmentalization, increased diffusion due to the enzymatic reaction,^62,63^ and multiphase reorganization/repartitioning. Specifically, we hypothesize that the pH-induced molecular reorganization of the multiphase coacervate system may contribute to the increased rate of reaction at early times, possibly by shepherding the reactants/products between the phases.

**Figure 6.**
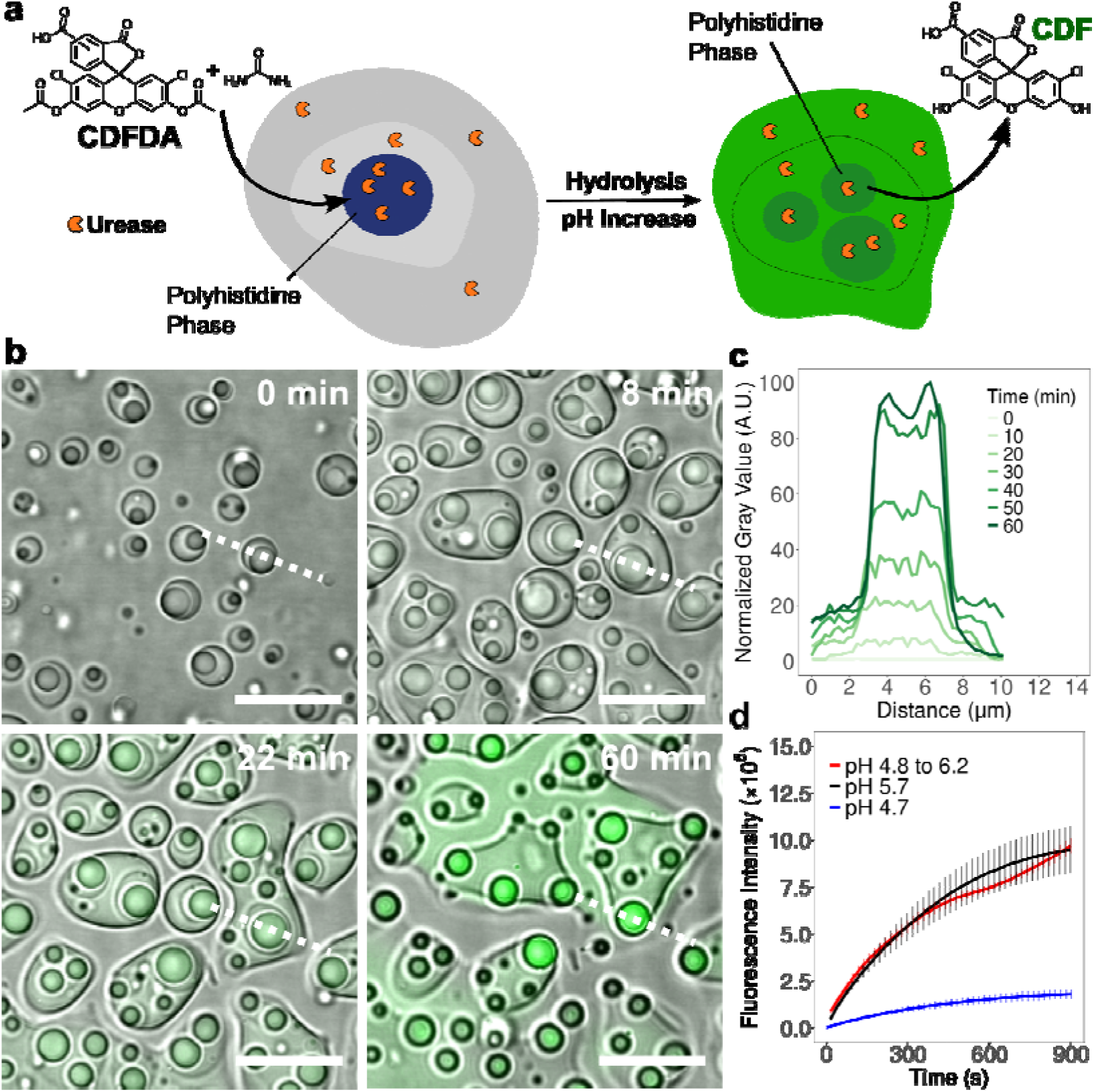
Kinetics of esterase-like catalysis in a changing pH environment. **a)** Schematic showing morphological changes and distribution of fluorescent product as the pH increases during the pH change and CDFDA hydrolysis in the pAsp multiphase. **b)** Merged confocal microscopy images of the transmitted light (DIC, greyscale) and CDF fluorescence (shown in green) channels showing structural changes in the coacervates as the pH increases and the hydrolysis of CDFDA into the green fluorescent dye (CDF) over a period of 60 min for the pAsp multiphase. **c)** Line profile plots showing the increase in fluorescence intensity across the multiphase over the course of the 60 min reaction (plots correspond to dashed lines in (**b**). **d)** Fluorescence spectroscopy kinetics showing the hydrolysis of CDFDA in the pAsp multiphase at static pH’s of 4.7 and 5.7 and in a pH changing system from pH 4.8 to 6.2 over the course of 15 min. Error bars in **d** represent standard deviations of a minimum of 3 experiments under each set of conditions. Scale bars are 10 µm.

The reaction rate at a static pH of 4.7 (blue line) was found to be approximately 7× slower than both the enzyme-free system at pH 5.7 (black line) and, most interestingly, the rate of reaction during the pH change reaction using urease had the fastest rate of all, approximately 1.2× faster than the static system at pH 5.7 (red line, Fig. 6d). Furthermore, the hydrolysis reaction rate as urease increases the pH decreases at approximately pH 5.8 before increasing further to form a reverse s-shaped kinetic plot (Supp. Fig. 30). This pH corresponds approximately to the point at which the DEAE-dextran-rich phase appears when the pH is increased from 4.7 to 6 in earlier experiments (Fig. 4 and Supplementary Video 1). This pH most likely corresponds to the approximate pKa of the pHis imidazole moieties in this system.

## Conclusions

This work demonstrated that aggregation-prone hydrophobic polymers such as pHis can be stabilized against aggregation by incorporating them into a multiphase coacervate system. This is especially important as transition metal catalysts and active site groups in native enzymes tend to be hydrophobic in nature.^64^ In native enzymes, the active site is stabilized in solution by the surrounding protein backbone. In our system, stabilization is instead achieved through multiple disordered coacervate phases, potentially offering a new strategy for supporting hydrophobic catalytic sites. In addition, by integrating a pH-sensitive polymer like pHis we achieved pH-dependent reorganization of phases within the coacervate droplets near biological pH. These morphological and microenvironment changes were influenced by the secondary structure of pHis and could be altered by the choice of polyanion (ATP or pAsp). The pH of the system was controllable using urease or GOx enzymes, and the resulting morphological changes were fully reversible. Importantly, we leveraged the inherent catalytic activity of pHis to achieve esterase-like activity in an enzyme-free multiphase system, with enhanced catalytic performance driven by pH-mediated phase perturbations. This rearrangement promotes polymer movement between phases, further boosting reaction rates. Our findings build on prior work showing that reaction rates in aqueous two-phase systems can be enhanced by separating products and starting materials into different phases.^65,66^ However, unlike those systems, our approach allows for dynamic modulation of polymer arrangement and phase composition through external stimuli, such as pH changes, to achieve greater rate enhancement.

Recent studies have shown that short peptides can form catalytic coacervates either by maintaining non-equilibrium droplet states through internal reactions, such as an aldol condensation reaction catalyzed by β-alanine amines,^67^ or by organizing flexible peptides into catalytically competent folded domains within self-coacervated droplets.^68^ These systems demonstrate the power of single component peptide-based compartments for achieving selective catalysis and structural regulation. Our findings build on this work by showing how synthetic, multiphase systems can also achieve stimulus responsive catalysis while maintaining compatibility with diverse polymer chemistries. Unlike single-phase or single component systems, our approach allows for dynamic modulation of polymer arrangement and phase composition through external triggers such as pH changes, enabling adaptive enhancement of catalytic rates.

A key feature of our system is its use of histidine, a residue frequently found in enzymatic active sites including many that contain catalytic triads, where each amino acid is positioned with nanometer-level precision. Our research suggests that clustering catalytic groups at high, localized concentrations in a disordered system can still result in effective catalytic activity. Given the abundance of ester groups in synthetic polymer environmental pollutants,^69^ histidine-based multiphase systems could be used to degrade such chemicals in an eco-friendly way. The next step in achieving this goal would be to modify systems like ours to catalyze the hydrolysis of non-biological molecules, which is a challenging task. In the long term, it may be possible to expand on the concept of histidine-based catalytic coacervates by incorporating additional reactive functional groups. This could further enhance catalytic efficiency or adjust substrate specificity, enabling the targeted breakdown of other compounds, such as peptides ethers, or phosphate-based nerve agents.

## Acknowledgements

This work was primarily supported by the National Science Foundation (NSF grant no. EF-1935059), with DLS and CD supported by NSF grant no. MCB-2317529, and microRaman supported by NASA Exobiology grant no. 80NSSC22K0553. We acknowledge the Huck Institutes’ X Ray Crystallography Core Facility (RRID:SCR_024464) for use of the Wyatt Dynapro Nanostar DLS and Jasco J-1500 CD Spectrophotometer, and Julia Fecko for helpful discussions on sample preparation, as well as the Penn State Materials Characterization Lab for the use of the Horiba LabRam HR Evolution and Maxwell Wetherington for helpful discussions on microRaman sample preparation and method development.

