## Supplementary Information for "Dynamic and Catalytic Multiphase Coacervates"

### Supplementary Methods

#### Materials and Methods

All materials were used as received. Poly-L-histidine (pHis, average molecular weight 5,000-25,000 Da), diethylaminoethyl-dextran-fluorescein isothiocyanate (DEAE-dextran-FITC, average molecular weight 40,000 Da), sodium hydroxide, adenosine 5’-triphosphate disodium salt hydrate (ATP), adenosine 5’-diphosphate sodium salt (ADP), adenosine monophosphate (AMP), adenosine, 4-morpholineethanesulfonic acid (MES), acetic acid, sodium acetate, glucose, urea, dimethyl sulfoxide, *N*-(3-Dimethylaminopropyl)-*N*′-ethylcarbodiimide hydrochloride (EDC), 5(6)-Carboxy-2′,7′-dichlorofluorescein diacetate (CDFDA), 5(6)-Carboxy-2′,7′-dichlorofluorescein (CDF), sodium cyanoborohydride, glucose oxidase (≥ 100,000 units/g) and urease (50,000-100,000 units/g) were purchased from Sigma-Aldrich. Poly-L-lysine hydrochloride, (pLys, PDI = 1.00-1.20, average molecular weight 16,000 Da) and poly-L-aspartic acid (pAsp, 1.00-1.20, average molecular weight 11,500 Da) were purchased from Alamanda Polymers. Alexa Fluor 633 maleimide, Alexa Fluor 488 hydrazide, rhodamine NHS, silicon spacers (Press-to-Seal Silicone Isolator, with 20 mm internal diameter and 0.5 mm depth) and Pierce™ Protein Concentrators 3K and 5K molecular weight cut-offs (MWCO) (both ≤ 0.5 mL volume) and Nalgene Rapid-Flow PES filter (0.22 µm) were purchased from ThermoFisher Scientific. Carboxymethyl-dextran (CM-dextran) was purchased from TdB Labs (average molecular weight 40,000-60,000 Da). A custom peptide consisting of 20 histidine residues terminated by a cysteine residue on the C-terminus was purchased from GenScript (pHis20-C). Hydrochloric acid, ethanol, toluene, and micro cover glass slides for confocal experiments (no. 1.5, 24 × 30 mm) were purchased from VWR. of *N*-(triethoxysilylpropyl)-o-polyethylene oxide urethane was purchased from Gelest. Bio-Gel P-6 fine polyacrylamide beads for size exclusion chromatography (100g, 45-90 µm, 1,000-6,000 MW fractionation range) were purchased from Bio-Rad Laboratories, Inc. Cover slides for microRaman (beveled edge, clipped corners, 25$\times$75$\times$1 mm, frosted 1 end, 1 side) were purchased from Globe Scientific Inc. Glass capillaries (Rectangle Boro Tubing 3520-050 Material / Length: Borosilicate / 050 mm) were purchased from VitroCom. All solutions were prepared using Milli-Q ultrapure water (18.2 MΩ·cm resistivity at 25 °C).

#### Fluorescent Labeling of Polyelectrolytes and Enzymes

Fluorescent variants of polyelectrolytes were synthesized using commercially available dyes via conjugation chemistry¹. Labeled pLys was prepared using NHS ester chemistry: 1.2 mol equiv of dye per polymer molecule was added to a 5 mg·mL⁻¹ solution of pLys in HEPES buffer (20 mM, pH 7.4). The reaction was stirred overnight and purified using a size-exclusion column and concentrated with a centrifugal filter (MWCO 5 kDa). pAsp and CM-dextran were labeled using hydrazide dyes. A 25× molar excess of EDC per polymer molecule was added to a 5 mg·mL⁻¹ polymer solution in MilliQ water (adjusted to pH 5 with 1 mM HCl). After ~15 min, 1.2 equiv of hydrazide dye (for pAsp) or 5 equiv (for CM-dextran) were added, and the reaction was stirred overnight. The next morning, 5 equiv of sodium cyanoborohydride were added to stabilize the hydrazone linkage. The labeled polymers were purified by size-exclusion chromatography and concentrated using centrifugal filters (MWCO 5 kDa). Instead of directly labeling pHis, a custom peptide consisting of 20 histidine residues with a C-terminal cysteine (pHis20-C) was used. Standard maleimide chemistry was used to label the Cys-terminated peptide. A 1.2 molar equivalent of maleimide dye was added to a 5 mg·mL⁻¹ solution of the peptide in HEPES buffer (20 mM, pH 7.4). The labeled peptide was purified by dialysis (MWCO 1 kDa) and concentrated using centrifugal filters (MWCO 1 kDa). Urease and glucose oxidase were also labelled using the same NHS ester chemistry described above for pLys. All molecules were lyophilized prior to use and were stable for ~1 month at 4 °C, or were stored at –22 °C for long-term storage.

#### Molecular Weights of Polyelectrolytes Used to Calculate Concentrations

The molecular weight of the monomer was used when calculating the molecular weights of each polymer. The molecular weight of peptides was calculated using the amino acid molecular weight less one water as an approximation for the charge concentration (pHis = 137.15 Da, pAsp = 115.09 Da, pLys = 128.18 Da, all molecular weights are per monomer). The molecular weight used when making stock solutions for coacervation was always the monomer molecular weight and not the molecular weight of the full polymer. Glucose was used as the monomer molecular weight of DEAE-dextran and CM-dextran (180.16 Da) and the full molecular weight of ATP disodium salt hydrate was used (551.14 Da) for ATP.

#### Formation of Coacervates

All polyelectrolyte stock solutions were prepared to a final concentration of 50 mM monomer concentration in Milli-Q water. pHis was dissolved by adjusting the pH to 4.5-4.8 using 1 mM HCl (measured with Mettler Toledo pH sensor InLab Ultra Micro-ISM pH probe). Multiphase coacervates were generated by first preparing single-phase coacervates at a final monomer charge concentration of 10 mM for each polyelectrolyte and a molecular concentration of 10 mM for ATP. “Monomer charge concentration” refers to the molar concentration of ionic charges along the polymer backbone (e.g., pAsp contributes one negative charge per monomer unit, so 10 mM monomer charge concentration corresponds to 10 mM in monomer units). In contrast, for ATP the concentration is expressed per molecule, irrespective of its multiple charges. The resulting single-phase coacervates were then combined to produce multiphase droplets. The three single-phase coacervates were made in the following order: pHis/ATP, pLys/pAsp, and DEAE-dextran/CM-dextran. pHis/ATP was prepared in acetate buffer (10 mM, pH 4.5), pLys/pAsp and DEAE-dextran/CM-dextran were both prepared in MES buffer (10 mM pH 6). When mixed together, this combination gave a final pH of 5.7. The different pH buffers were required to increase solubility of pHis, while allowing a final pH after mixing of approximately pH 5.7. To achieve a final pH of 4.7, the three single-phase coacervates were all prepared in acetate buffer (10 mM, pH 4.5) before mixing. When making the single-phase coacervates, the cation was always added last to avoid the polycation irreversibly sticking to the surface of the plastic microcentrifuge tubes. Subsequent experiments on single-phase and multiphase systems were carried out within 1hr on coacervates prepared in this manner without noticeable degradation. The pH values of 4.7 and 5.7 were measured for the base coacervate system and are expected to apply to subsequent experiments, although minor variations may occur between preparations.

#### PEGylation of Coverslips and Capillaries

Glass coverslips or capillaries were soaked for 30 min in a solution of ~1 M KOH in isopropanol, after which they were washed thoroughly with ethanol followed by Milli-Q water and dried in the oven at 60 °C overnight. Approximately 900 mg of *N*-(triethoxysilylpropyl)-o-polyethylene oxide urethane was dissolved in 300 mL of toluene. The washed glass was submersed in the toluene solution and left to react for 4 hr. The toluene solution was removed, and the glass was washed thoroughly with ethanol and then MilliQ water before being left in the oven to dry overnight at 60 °C before use.^2^

#### Confocal Microscopy and Image Analysis

Confocal images were acquired using a Leica TCS SP5 laser scanning confocal inverted microscope. Samples were prepared as described in the Formation of Coacervates section, with the addition of a small amount of fluorescently labeled polymer or its equivalent (pHis20-C) prior to the final mixing step to form the multiphase. The amount of labeled fluorophore added was adjusted for each system to ensure sufficient signal intensity while minimizing any potential effects on multiphase stability. Samples were imaged on custom-made poly(ethylene glycol)-functionalized coverslips to prevent coacervate wetting on the glass surface. All samples were prepared and imaged promptly after preparation. Fluorescence intensity values were obtained from raw images in Fiji following background correction. Droplets were counted using the Particle Analysis plugin in Fiji, with a circularity threshold of 0.5.

#### Quantification of Relative Fluorescent Intensities

Two normalization strategies were applied depending on the type of plot. For bar plots displaying a single fluorophore (one graph per dye), intensities were normalized to the maximum average intensity observed across all phases for that dye; this maximum was set to 100%, and all other values were scaled relative to it. This approach emphasizes the relative distribution of each fluorophore across coacervate phases. For plots combining all fluorophores and phases on a single graph, intensities were normalized to the maximum observed intensity across all fluorophores and phases (typically Alexa Fluor 633). This allowed direct comparison of fluorophore enrichment across phases, although weaker fluorophores may appear underrepresented when compared to the brightest dye. In all cases, normalization was based on ≥20 intensity measurements per phase, and error bars represent standard deviations. As fluorescence is influenced by the physicochemical properties of the local coacervate phase, we avoided using environment-sensitive rotor dyes;^3^ however, phase-dependent intensity effects cannot be fully excluded.

#### Raman Spectra and Raman Microscopy

Raman spectra were collected using a Horiba LabRam HR Evolution spectrometer equipped with a 532 nm laser line and a Freespace Olympus BX51 upright confocal microscope. Multiphase coacervates were prepared as described in the *Formation of Coacervates* section above at pH 5.7. For imaging with the upright microscope, samples were prepared by suspending coacervate droplets from the upper interior surface of silanized capillaries, which were loaded by horizontally dipping the tip of the capillary into the coacervate suspension. The capillaries were mounted onto supporting coverslips and sealed at both ends with Scotch tape. Droplets were allowed to equilibrate for at least 30 minutes before imaging. Imaging was performed using a Zeiss 100X oil immersion lens with the 532 nm laser operating at 100% power. To determine the optimal focal plane for droplet microRaman imaging, a Z-depth profile was acquired in 1 µm steps with 2 seconds of accumulation per step. The focal plane was selected based on the Z-depth that yielded the greatest integrated intensity in the C–H stretching region (2800–3182 cm⁻¹). After selecting the appropriate focal plane, a Swift map was acquired with 0.99 seconds of accumulation per pixel to generate Raman images of the droplets.

#### Quantification of Relative Raman Intensities

To analyze ATP partitioning, we quantified the 719 cm⁻¹ Raman band corresponding to phosphate stretching vibrations.^4-6^ MicroRaman maps were converted to grayscale and analyzed in Fiji. For each droplet, a line was drawn across the image from the inner coacervate phase to the surrounding dilute phase, and average signal intensities were extracted for each phase along the line scan. Intensities were then normalized by setting the sum of signals from all phases within the droplet to 100%, and expressing the signal from each phase as a fraction of this total. This procedure was repeated for all imaged droplets (n > 10).

#### Circular Dichroism Spectroscopy

Poly-L histidine (50 mM monomer concentration) was dissolved in Mili-Q water and pH adjusted to ≈ pH 4 (verified Supelco pH-indicator strips pH 2.0-9.0) with 2 M HCl to fully solubilize. Stocks were then pH adjusted to 4.5 and 6.0 with NaOH for analysis (measured by pH probe). CD spectra were collected using a Jasco J-1500 CD Spectrophotometer, scanning at 50 nm/min with a 4-second data integration time in a 0.1mm cuvette. Samples were collected in triplicate with 3 acquisitions per sample. Samples were prepared by diluting pHis at the appropriate pH in Mili-Q water. pH test strips were used to verify pH after dilution. For samples containing ATP/pAsp, ATP and pAsp stocks were individually pH adjusted to 4.5 and 6.0 prior to use. pHis was diluted in the appropriate amount of water followed by addition of ATP/pAsp at the correct pH. Data was normalized to the mean residue ellipticity by the formula

$$\frac{MRW\times\theta}{10\times d\times c}$$

Where MRW = mean residue weight = 137.146 Da = MW of histidine – 18.0 Da (water) because only histidine residues are present, $\theta$ = observed ellipticity (milidegrees), d = pathlength (cm), c = concentration (mg.mL^-1^) to give the final units of deg × cm^2^ × dmol^-1^.

#### Dynamic Light Scattering

Dynamic light scattering (DLS) data were collected on a Wyatt Dynapro Nanostar DLS detector. Mili-Q water, NaOH, and HCl used for stock and sample preparation were filtered through a 0.22 μm filter (Thermo Scientific Nalgene Rapid-Flow PES filter). pHis stocks were prepared in effort to avoid pH dependent pHis aggregation; pHis was dissolved in filtered Mili-Q water and pH adjusted to pH 4.0-4.5, verified with pH test strips. pHis, ATP, and pAsp stocks were filtered with 0.22μm syringe filters (Millipore) and the pH of each was adjusted to pH 4.5 or 6.0, verified via pH probe. Sample components were added in fixed order; pHis was diluted with an appropriate volume of water, followed by the addition of pAsp or ATP (except in no polyanion trials). Samples were gently mixed and immediately transferred to cuvettes for measurement.

#### Fluorescence Recovery After Photobleaching (FRAP)

FRAP experiments were performed on pHis-rich phases doped with fluorescently labeled pHis20-C peptide conjugated to Alexa Fluor 633. We used the same image settings as for the previous experiments. A 10-frame pre-bleach sequence was followed by a 10-frame bleach at 100% 633 nm, 100% 543 nm, and 10% 488 nm laser powers. Regions of interest were 1 μm in diameter, and droplets were imaged with a 1400 Hz scanning speed. For accurate comparison across droplets, we selected coacervates that were approximately the same size and, where possible, ensured that the droplet diameter was larger than the bleached region; however, this was challenging due to the small size of many multiphase coacervates. FRAP curves were fitted to a single exponential recovery function in R software based on the following equation:

𝑁(𝑡) = 𝐴 (1 − exp (−𝑡 /𝜏)) + 𝐶

Where A and C are constants, and 𝜏 is the fluorescence recovery time. The halftime of recovery,

𝑡_1/2_, is calculated using 𝜏_1/2_ = 𝜏ln (2).

#### Change in pH Using Glucose Oxidase and Urease Enzymes

Multiphase coacervates were prepared as described in the formation of coacervates section above with the extra addition of 0.2 mg.mL^-1^ of urease (when increasing the pH) or glucose oxidase (when decreasing the pH). The desired enzyme was added to the final solution while keeping the concentrations of the other components constant. The reaction was initiated by the addition of 5 mM urea (urease) or 15 mM glucose (glucose oxidase) which were both dissolved in MilliQ water. The change in pH was measured using a pH Probe, taking a data point either every 30 s (urease) or 120 s (glucose oxidase) until the pH stabilized.

#### Imaging Change in pH Using Confocal Microscopy

Multiphase coacervates were prepared and doped with the relevant fluorescently labelled polyelectrolyte as described previously, with the addition of either glucose oxidase or urease enzymes (0.2 mg.mL^-1^). Labelled versions of the enzymes were not used in these experiments to preserve the stability and the kinetics of the enzymes. The coacervate solution was added to a PEGylated glass coverslip, the pH change reaction was initiated by addition of the relevant substrate (5 mM urea or 15 mM glucose) to the outer edge of the coacervate solution. Images were taken at a frequency of either every 30 s for experiments lasting 15 min or every 2 min for experiments lasting 60 min.

#### Histidine-mediated Hydrolysis of CDFDA in Bulk Solution

Hydrolysis kinetics were collected using a Horiba FluoroLog-3 spectrofluorometer equipped with a Peltier-based temperature control which was kept at 25 °C for all experiments. Multiphase coacervates were prepared using the same procedure as described above in the *Formation of Coacervates* section. 5(6)-Carboxy-2′,7′-dichlorofluorescein diacetate (CDFDA) was dissolved in DMSO to a concentration of 2 mM, this was subsequently diluted to a final concentration of 25 µM in Milli-Q water. This solution remained stable for approximately 5 hours before precipitating, likely due to its tendency to aggregate in Milli-Q water. To initiate the reaction, first the multiphase coacervate was added to a cuvette and left to equilibrate to 25 °C for 2 min. After, CDFDA (25 µM) was added to a final dye concentration of 5 µM. A data point was recorded every 2 s for a total of 900 s, using a λ_ex_ of 492 nm and a λ_em_ of 517 nm. The experiments were repeated in triplicate and averages and standard deviations were taken. The different pH’s were obtained in the same manner as described in the coacervate preparation methodology. Single-phase coacervates were recorded by not adding the other two single phases and instead adding an equal volume of buffer. For the pH 5.7 system, a 3:1 ratio of MES buffer (10 mM, pH 6) to acetate buffer (10 mM, pH 4.5) was used. For the pH 4.7 system, only acetate buffer (10 mM, pH 4.5) was used. pH change experiments were carried out by adding urease (0.2 mg.mL^-1^) to the coacervates, the reaction was initiated using a solution of urea (25 mM) mixed with CDFDA (25 µM) to final urea and CDFDA concentrations of 5 mM and 5 µM respectively.

#### Small Volume Histidine-mediated Hydrolysis of CDFDA

Coacervates were prepared as described in the *Formation of Coacervates* section above. For the multiphase system, only pHis was fluorescently labeled by doping with Alexa Fluor 633-labelled pHis20-C peptide, to avoid spectral overlap with the excitation and emission of CDFDA. The reaction was initiated by adding CDFDA to a final concentration of 20 µM. The hydrolysis was followed using the 488 nm laser, taking images every 30 s for 15 min. Different pH’s were obtained by mixing acetate and MES buffer when preparing the coacervates, as described previously. pH change experiments were carried out by adding urease (0.2 mg.mL^-1^) to the coacervates, the reaction was initiated using a mixture of urea (25 mM) and CDFDA (25 µM) to final urea and CDFDA concentrations of 5 mM and 5 µM respectively, and the reaction was imaged every 30s for 1 hr.

### Supplementary Figures

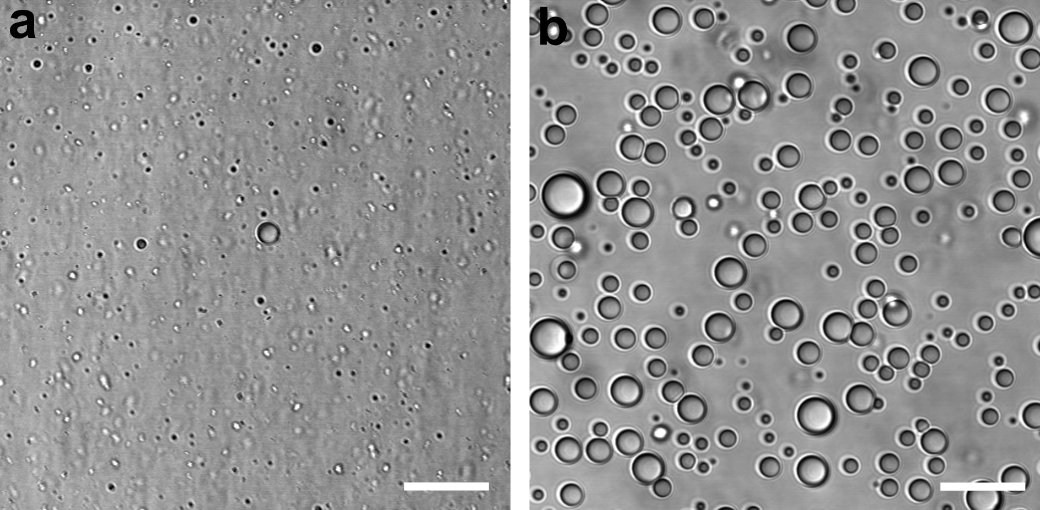

**Supplementary Figure 1. (a)** Brightfield images of pHis coacervates showing changes in droplet diameter at pH 6, when paired with pAsp **(a)** or ATP **(b)**. All polymers were used at a final monomer charge concentration of 10 mM; ATP was also used at a final molecular concentration of 10 mM. Scale bar: **(a)** = 20 µm, **(b)** = 10 µm.

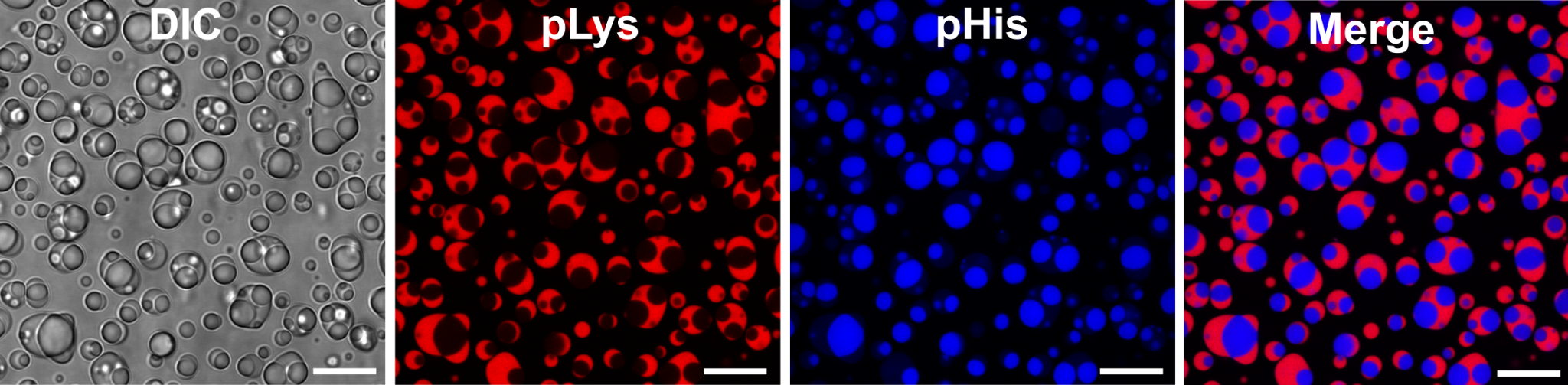

**Supplementary Figure 2.** Fluorescence confocal microscopy images of pHis/ATP and pLys/pAsp multiphase coacervates at pH 5.7. Single-phase coacervates were prepared separately before mixing together. Separate channels were false-colored for clarity. All scale bars = 10 µm.

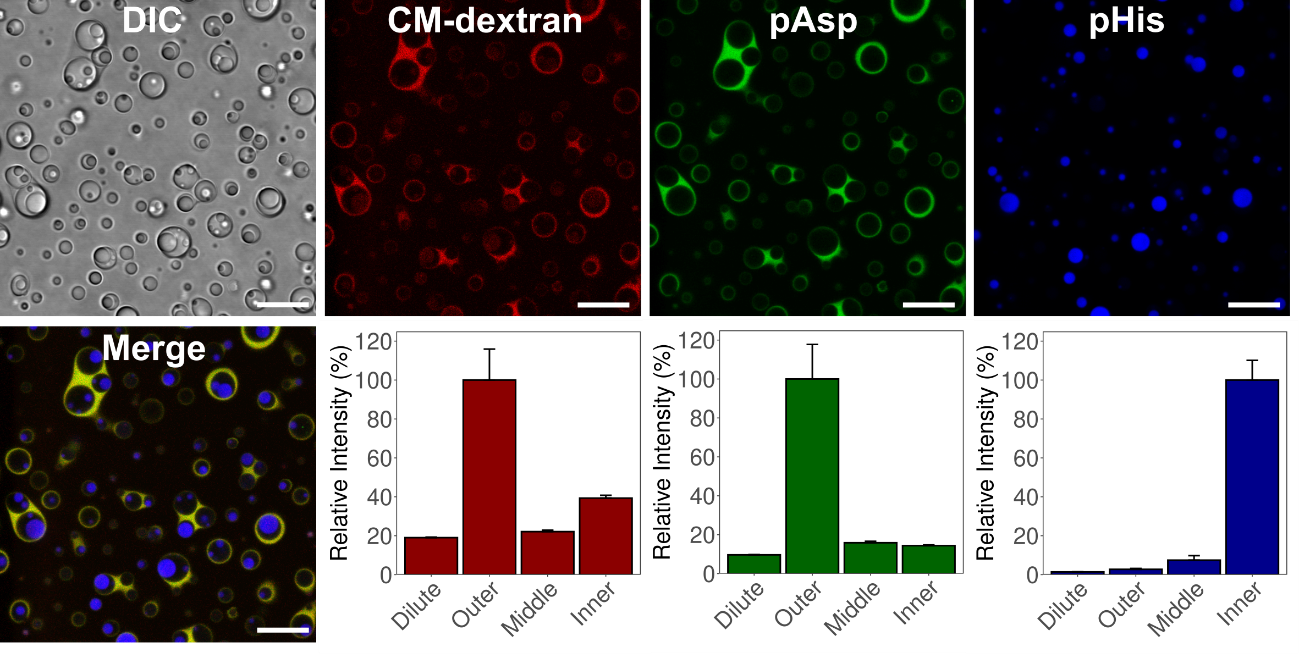

**Supplementary Figure 3.** Confocal microscopy images and corresponding relative intensity bar graphs showing the distribution of labeled polymers in ATP multiphase coacervates, highlighting the relative localization of polyanions and pHis at pH 5.7. The multiphase system was prepared by separately forming DEAE-dextran/CM-dextran, pLys/pAsp, and pHis/ATP coacervates prior to mixing them together. Labels in the figure correspond to the fluorescently labelled phases: CM-dextran was labeled with rhodamine, pAsp with Alexa Fluor 488, and pHis with Alexa Fluor 633. Bar plots display the relative fluorescence intensity in each phase, where the fluorescence intensities are normalized to the maximum signal for each dye, such that the phase with the highest intensity is set to 100%. Intensities in the other phases are expressed as a percentage relative to this maximum. Error bars represent standard deviations of a minimum of 20 droplets. Separate fluorescence channels were false-colored for clarity. Scale bars: 10 µm.

| **Peak (cm^-1^)** | **Assignment** |
| --- | --- |
| 719 | ATP: *ν_s_*(POP) triphosphate backbone stretching^4-6^ |
| 809 | pHis pH>6.5: out of plane bending of C-H, C-C in imidazole^7^ |
| 848 | ATP: adenine (skeletal mode, in-plane)^8^ |
| 878 | ATP: (major) and pHis pH=2.58 (minor): δ (N-9-H) out-of-plane^8^ |
| 989 | pHis: imidazole ring C-C rocking or scissoring: δ (R)^7^ |
| 1043 | pHis pH>6.5, side-chain skeletal stretching^9,10^ |
| 1108 | ATP (major) and pHis pH>2.58 (minor): *ν_e_* (PO_3_) O=P=O stretching, C-H in-plane bend^9^ |
| 1203 | pHis pH=2.58 and pHis>6.5: NCN sym. str. + N-H in-plane bend^9^ |
| 1257 | pHis: ring breath^9^ |
| 1299 | ATP and pHis pH>6.5^2^: ring breath^9^ |
| 1328 | ATP (major): *ν*(C-N), *ν*(C=N); pHis pH>6.5 (weak): δ (R)^8,12^ |
| 1368 | ATP: δ (C-8-H), δ (C-2-H) out-of-plane^8^ |
| 1434 | pHis methylene scissoring: δ (CH_2_)^10^ |
| 1474 | pHis pH=2.58 (major) and ATP (minor)5^,11^: N-H in-plane bend^8^ |
| 1494 | pHis pH=2.58 (major) and ATP (minor)^5,11^: N-H in-plane bend^8^ |
| 1568 | pHis pH=2.58 (major) and ATP (minor)^5,11^: C=C stretching^8^ |
| 1623 | pHis pH=2.58: C=C stretching^9^ |
| 1664 | pHis pH>6.5 (major) and pHis pH=2.58 (minor)^7,9^ |

### Supplementary Table 1.

Observed peaks and their respective assignments from microRaman microscopy experiments.

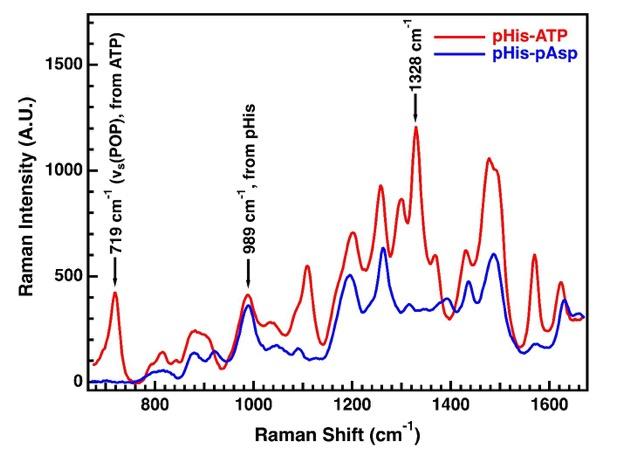

**Supplementary Figure 4.** Raman spectra of pHis/ATP (red) and pHis/pAsp (blue) single-phase coacervates, collected in MES buffer at pH 6. Peaks correspond to vibrational modes of the respective components in the condensed phase. For detailed peak assignments and experimental conditions, see *Supplementary Table 1* and the *Materials and Methods* section.

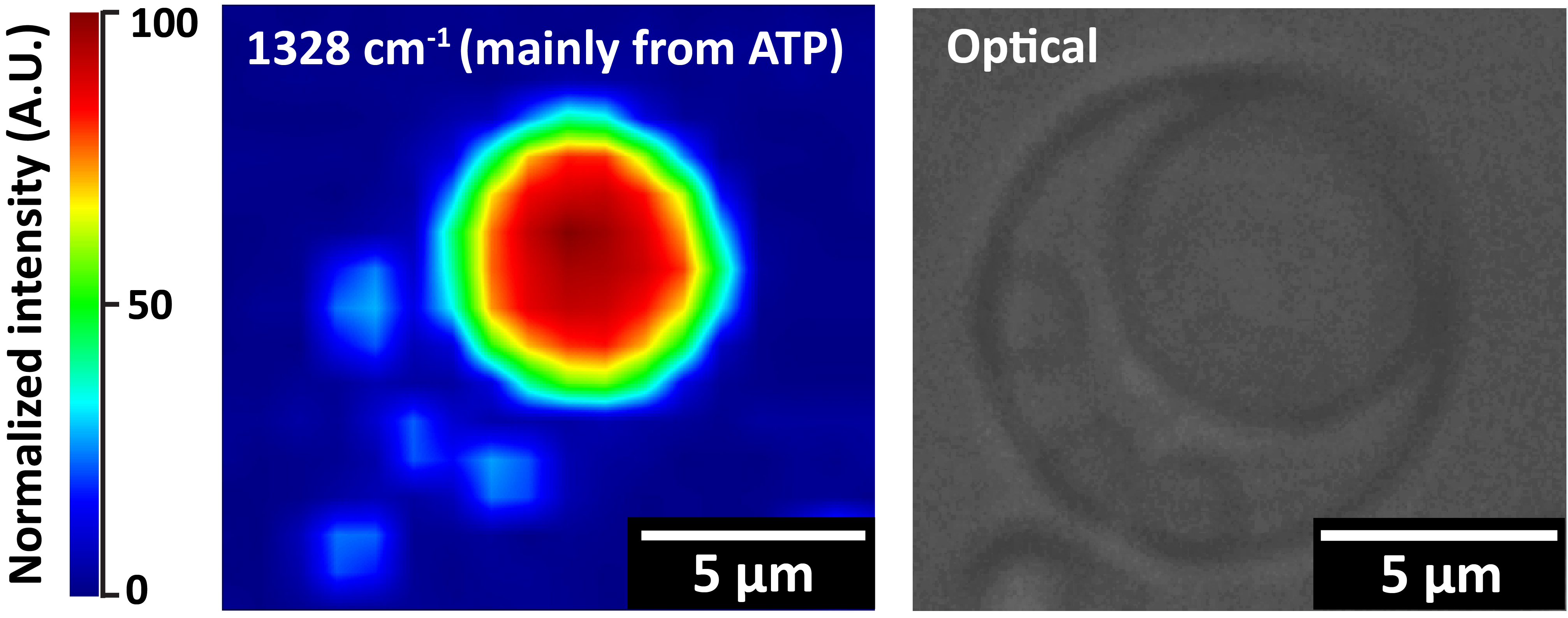

**Supplementary Figure 5.** microRaman map and corresponding optical image of a DEAE-dextran/CM-dextran, pLys/pAsp, and pHis/ATP multiphase coacervate droplet in MES buffer at pH 6. Raman intensities from the 719 cm⁻¹ phosphate stretching band of ATP were analyzed in Fiji and normalized by setting the total signal across all phases to 100%. For detailed experimental conditions, see the *Materials and Methods* section.

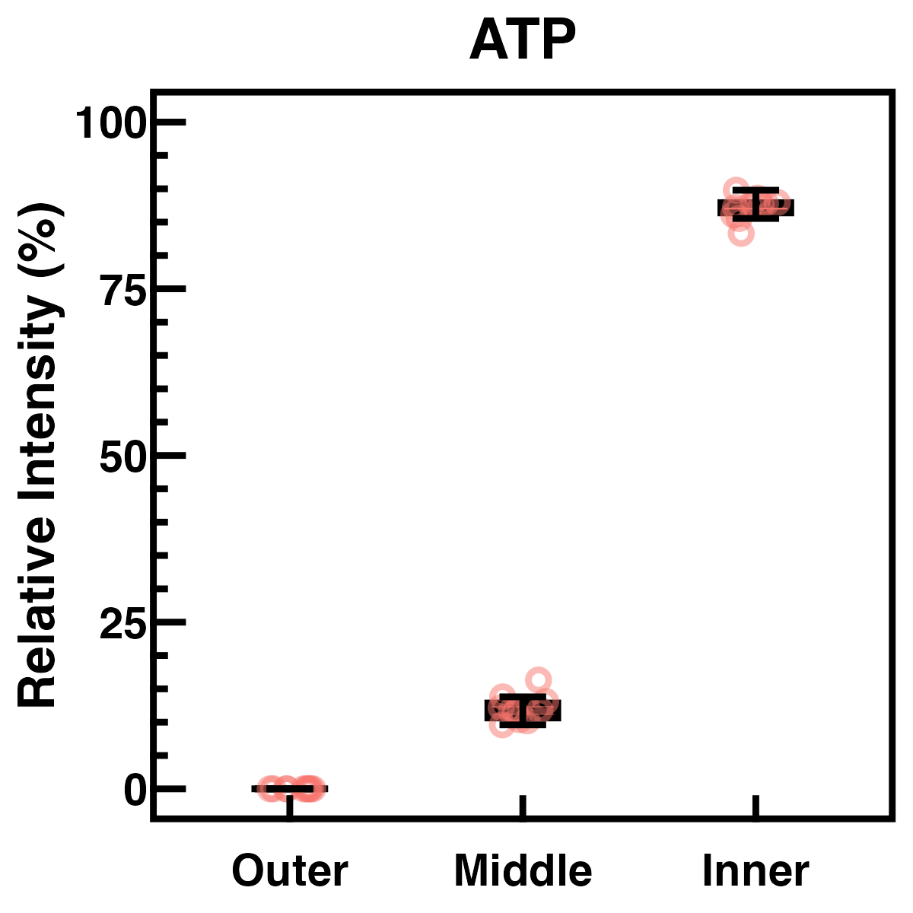

**Supplementary Figure 6.** Relative Raman intensity of ATP’s phosphate stretching ν_s_(P-O-P) at 719 cm^-1^, excited at 532nm laser. Raman intensities from the 719 cm⁻¹ phosphate stretching band of ATP were analyzed in Fiji and normalized by setting the total signal across all phases to 100%. Error bars represent standard deviations of 10 droplets among triplicates. Raman was not able to distinguish between the outer coacervate phase and the dilute phase (all denoted as outer phases) due to a low concentration of Raman-active functional groups in these phases.

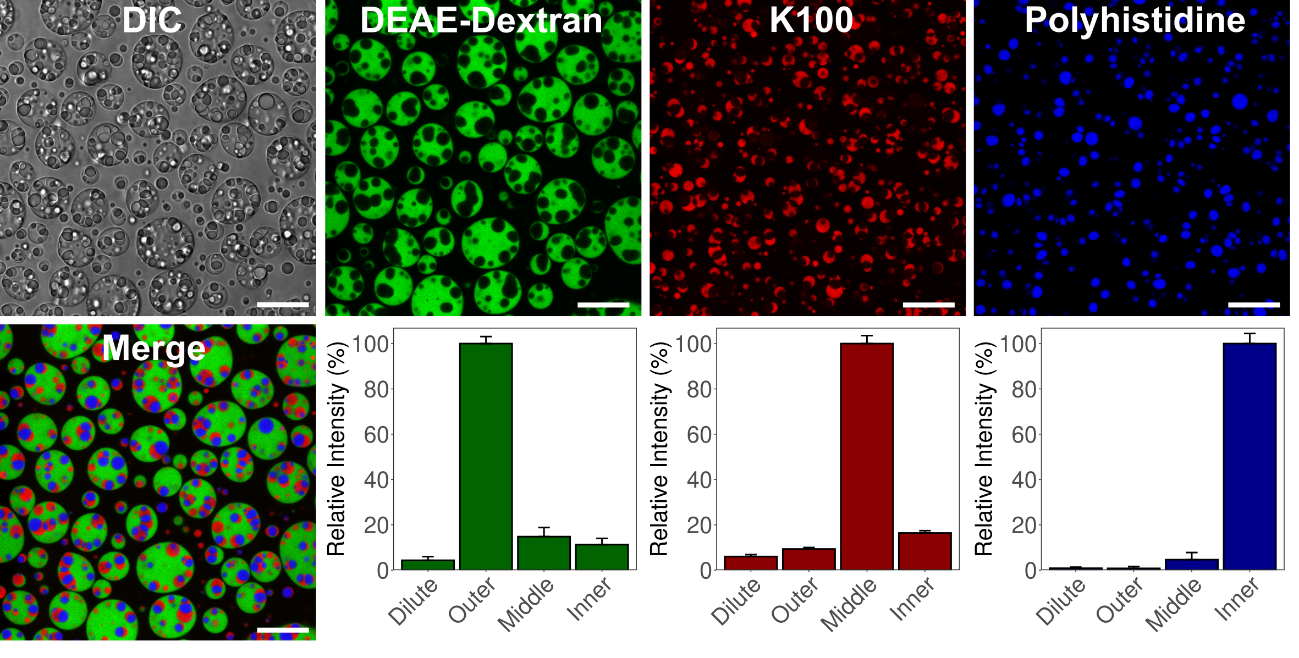

**Supplementary Figure 7.** Confocal microscopy images and corresponding relative intensity bar graphs showing the distribution of labeled polymers in multiphase coacervates, highlighting the relative localization of polycations and pHis at pH 4.7. The multiphase system was prepared by separately forming DEAE-dextran/CM-dextran, pLys/pAsp, and pHis/ATP coacervates prior to mixing them together. Labels in the figure correspond to the fluorescently labelled phases: DEAE-dextran was labeled with fluorescein, pLys with rhodamine, and pHis with Alexa Fluor 633. Bar plots display the relative fluorescence intensity in each phase, where the fluorescence intensities are normalized to the maximum signal for each dye, such that the phase with the highest intensity is set to 100%. Intensities in the other phases are expressed as a percentage relative to this maximum. Error bars represent standard deviations of a minimum of 20 droplets. Separate fluorescence channels were false-colored for clarity. Scale bars: 10 µm.

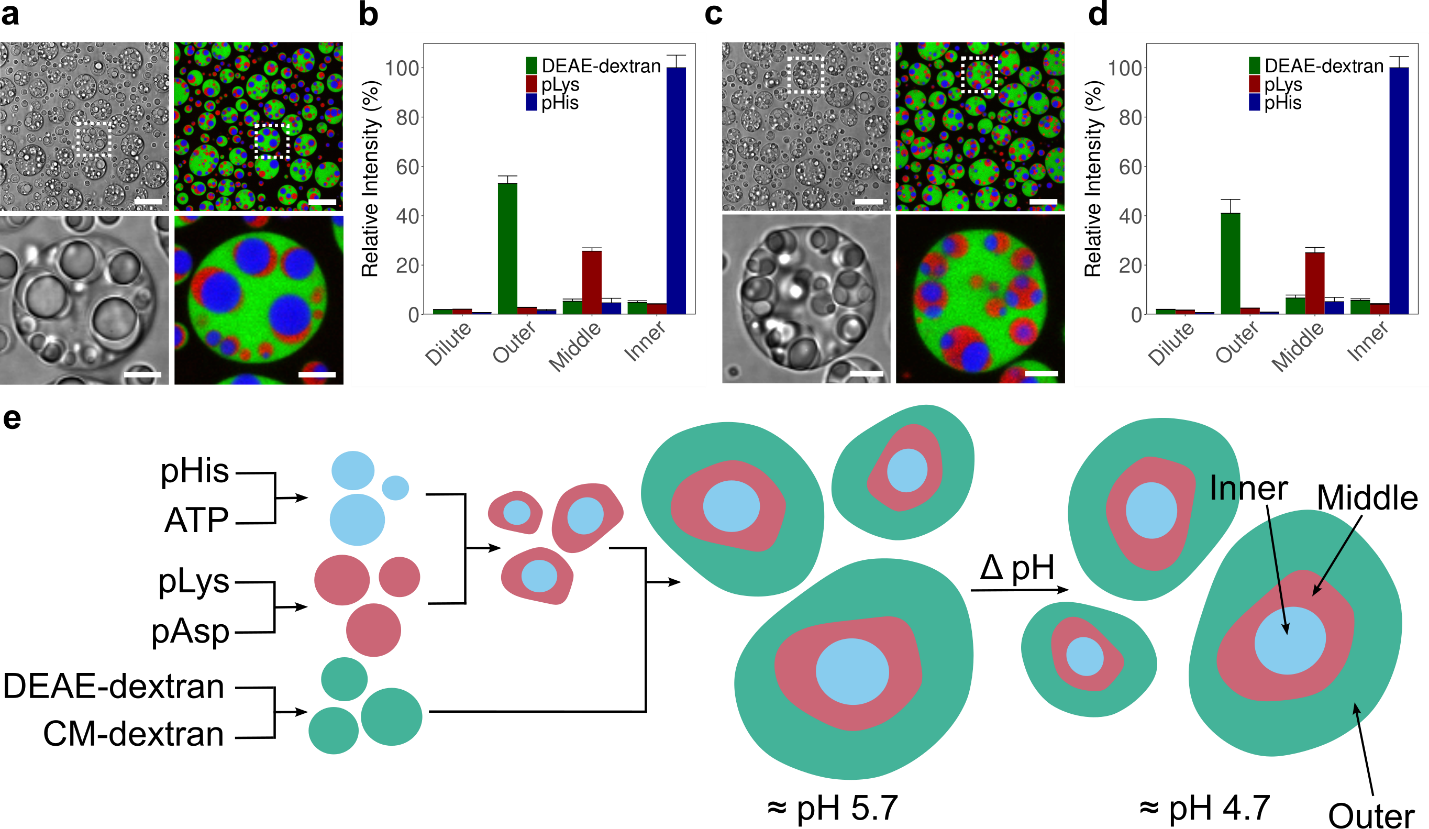

**Supplementary Figure 8 (a, c)** Confocal fluorescence microscopy images of DEAE-dextran/CM-dextran, pLys/pAsp, and pHis/ATP multiphase coacervates at pH 5.7 **(a)** and pH 4.7 **(c)**. **(b, d)** Corresponding bar plots showing the relative fluorescence intensities of DEAE-dextran-FITC (green), pLys-rhodamine (red), and pHis-Alexa Fluor 633 (blue) in the coacervate multiphase system at pH 5.7 (**b**) and pH 4.7 (**d**). Intensities are normalized such that 100% corresponds to the maximum intensity across all dyes and all phases, other dyes and phases are scaled relative to this value. A minimum of 20 measurements were collected per phase. Error bars represent standard deviations. Scale bars: 10 µm (main images), 5 µm (magnified images). **(e)** Schematic representation illustrating the pH stability of the system between approximately pH 5.7 and pH 4.7.

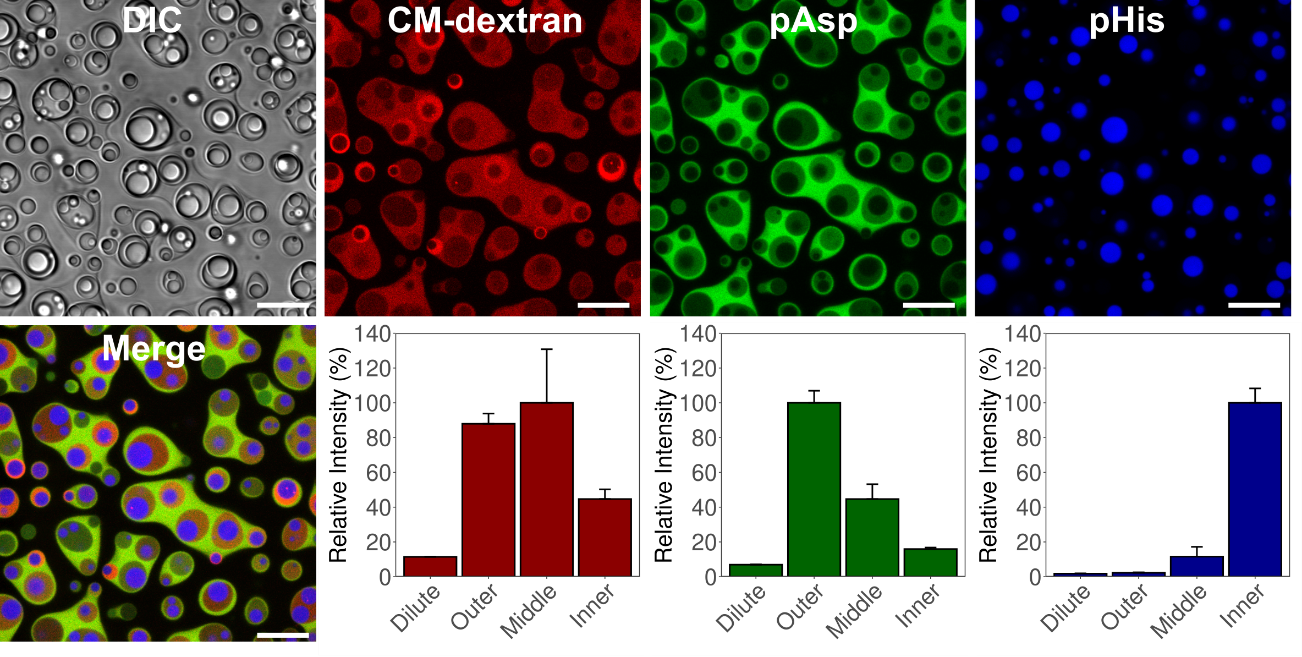

**Supplementary Figure 9.** Confocal microscopy images and corresponding relative intensity bar graphs showing the distribution of labeled polymers in multiphase coacervates, highlighting the relative localization of polyanions and pHis at pH 4.5 in the presence of ATP. The multiphase system was prepared by separately forming DEAE-dextran/CM-dextran, pLys/pAsp, and pHis/ATP coacervates prior to mixing them together. Labels in the figure correspond to the fluorescently labelled phases: CM-dextran was labeled with rhodamine, pAsp with Alexa Fluor 488, and pHis with Alexa Fluor 633. Bar plots display the relative fluorescence intensity in each phase, where the fluorescence intensities are normalized to the maximum signal for each dye, such that the phase with the highest intensity is set to 100%. Intensities in the other phases are expressed as a percentage relative to this maximum. Error bars represent standard deviations. Separate fluorescence channels were false-colored for clarity. Scale bars: 10 µm.

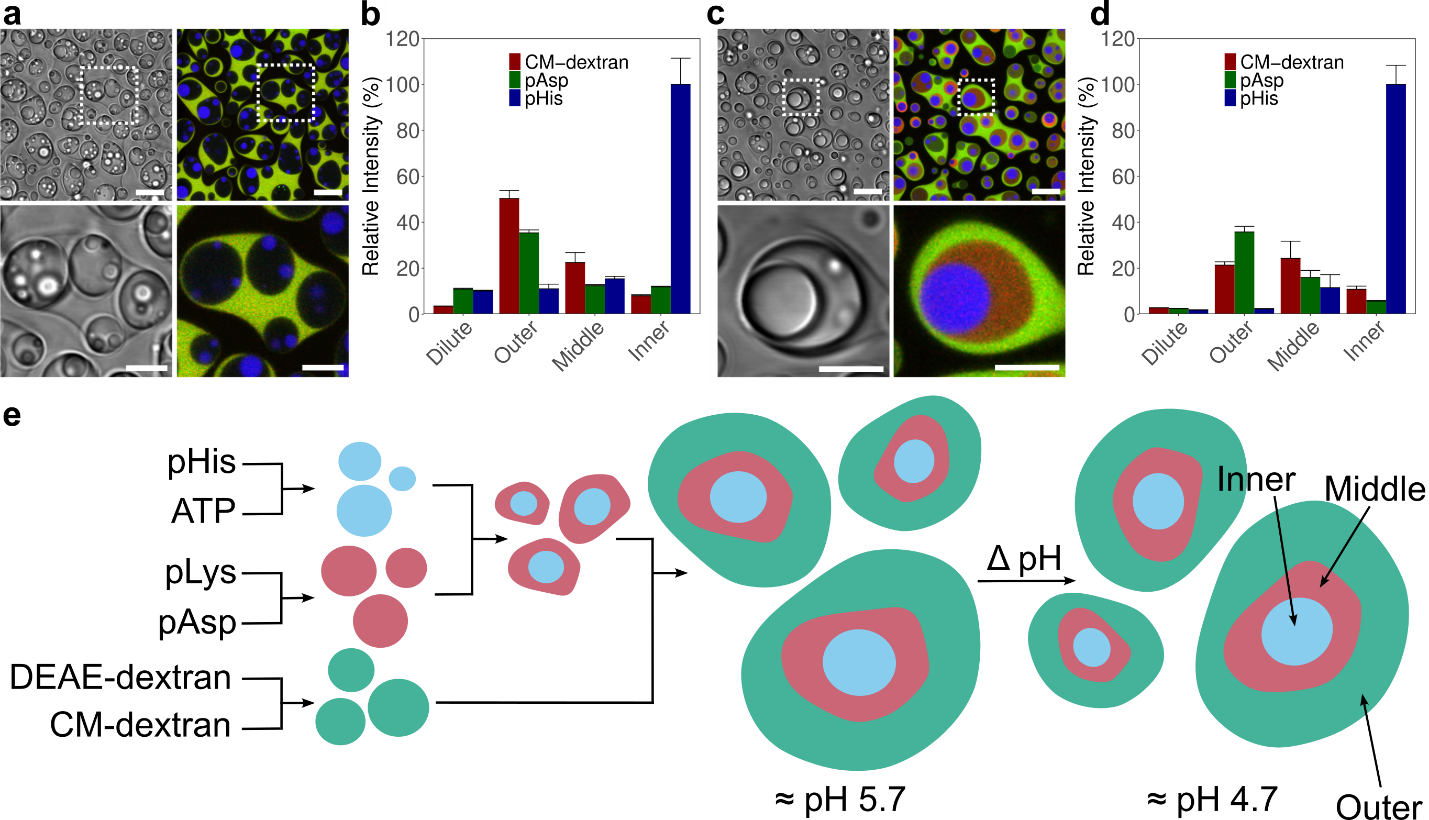

**Supplementary Figure 10 (a, c)** Confocal fluorescence microscopy images of DEAE-dextran/CM-dextran, pLys/pAsp, and pHis/ATP multiphase coacervates at pH 5.7 (a) and pH 4.7 **(c)**. **(b, d)** Corresponding bar plots showing the relative fluorescence intensities of CM-dextran-rhodamine (red), pAsp-Alexa Fluor 488 (green), and pHis-Alexa Fluor 633 (blue) in the coacervate multiphase system at pH 5.7 **(b)** and pH 4.7 **(d)**. Intensities are normalized such that 100% corresponds to the maximum intensity across all dyes and all phases, other dyes and phases are scaled relative to this value. A minimum of 20 measurements were collected per phase. Error bars represent standard deviations. Scale bars: 10 µm (main images), 5 µm (magnified regions). **(e)** Schematic representation illustrating the stability of the multiphase system between approximately pH 5.7 and pH 4.7.

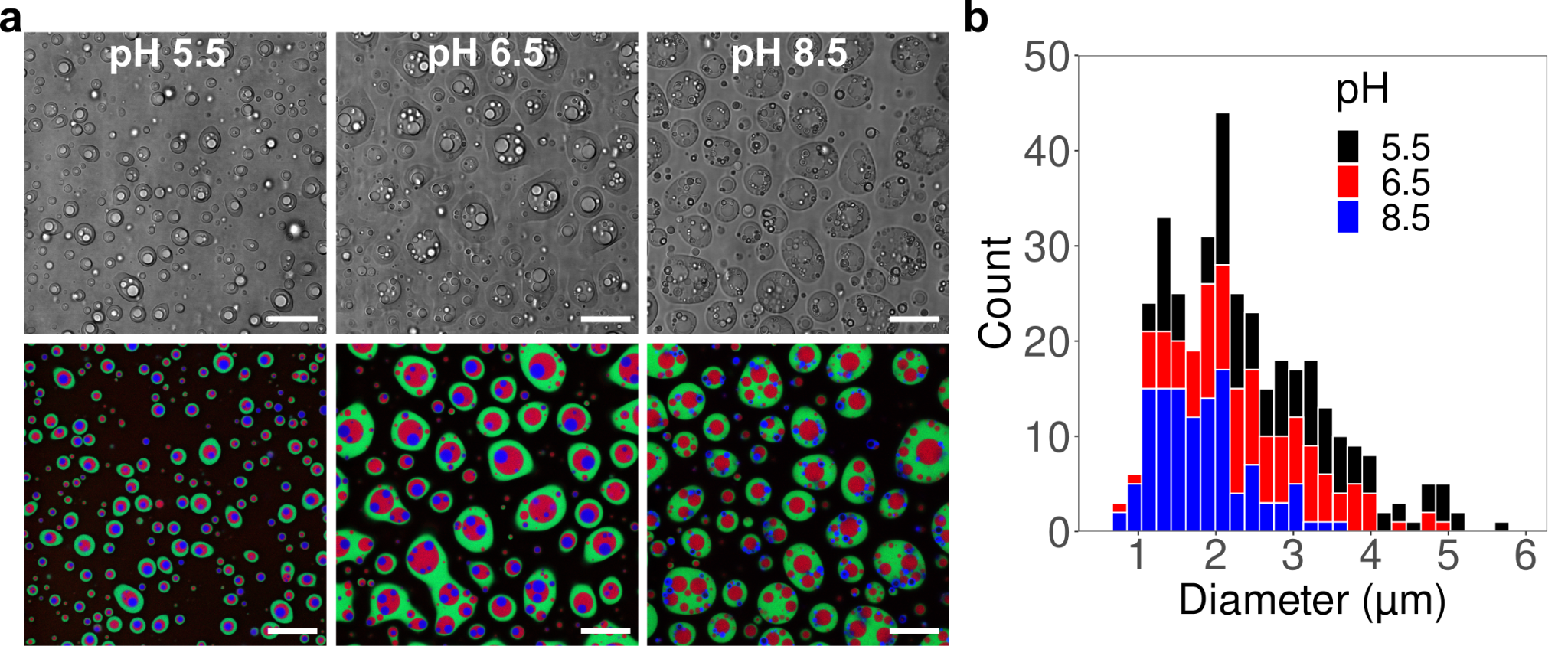

**Supplementary Figure 11.** Effect of pH on ATP-containing multiphase droplets. **(a)** Confocal microscopy images of DEAE-dextran/CM-dextran, pLys/pAsp, and pHis/ATP coacervates at pH 5.5, 6.5, and 8.5. DEAE-dextran was labeled with fluorescein (green), pLys with rhodamine (red), and pHis with Alexa Fluor 633 (blue). Separate fluorescence channels were false-colored for clarity. Scale bars: 10 µm. **(b)** Histograms of the diameters of the blue pHis-rich phase, based on 120 measurements at each pH. pH 5.5: mean = 2.79 µm, median = 2.69 µm; pH 6.5: mean = 2.48 µm, median = 2.35 µm; pH 8.5: mean = 1.81 µm, median = 1.74 µm.

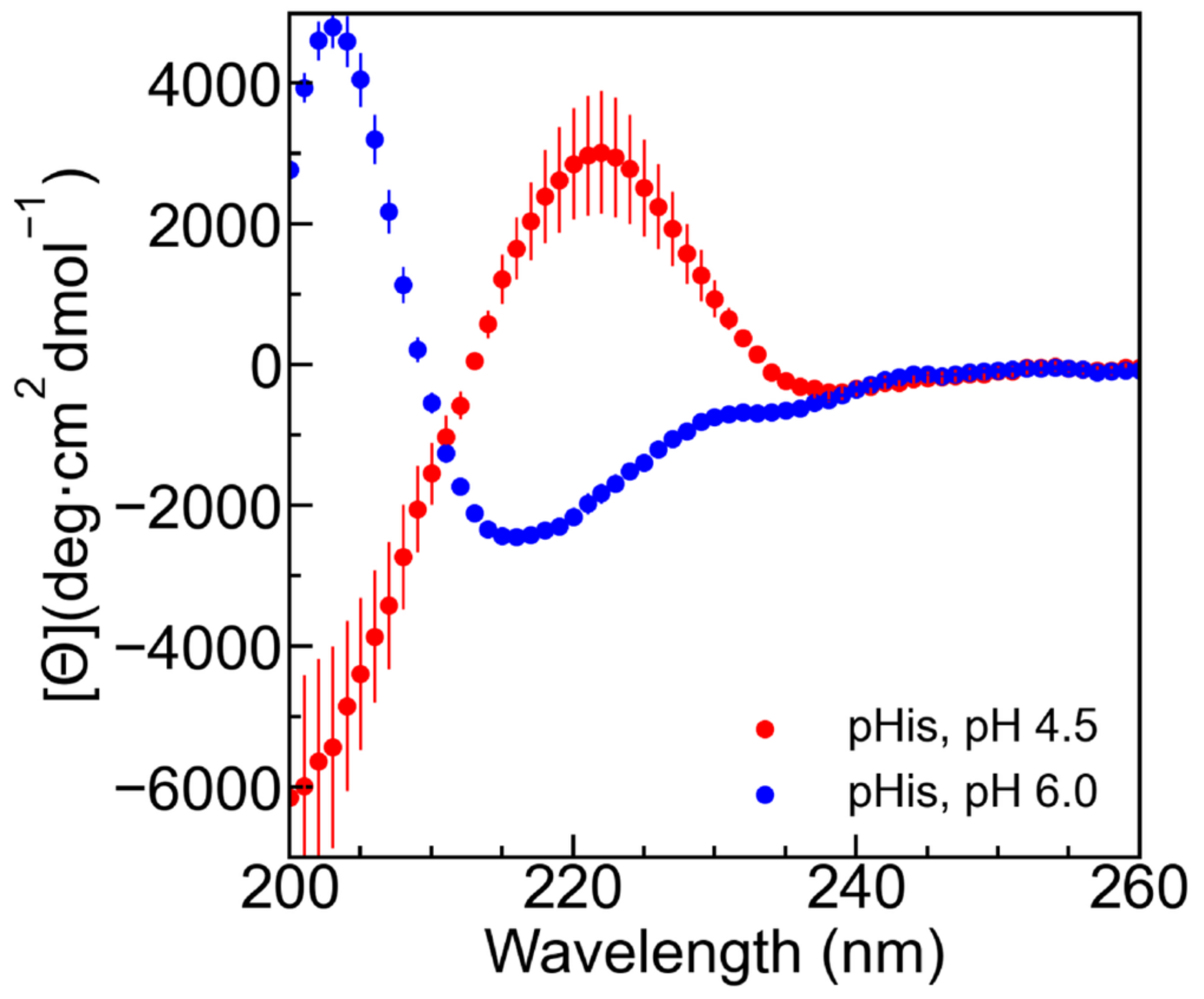

**Supplementary Figure 12.** Circular dichroism spectra of 2 mM monomer pHis at pH’s 4.5 and 6.0, showing coil and β-sheet structures respectively. CD signal normalized to mean residue ellipticity ([Θ]) as described in methods. Stock solutions were pH adjusted using dilute solutions of HCl and NaOH to achieve the desired pH in the absence of added buffer or added salts.

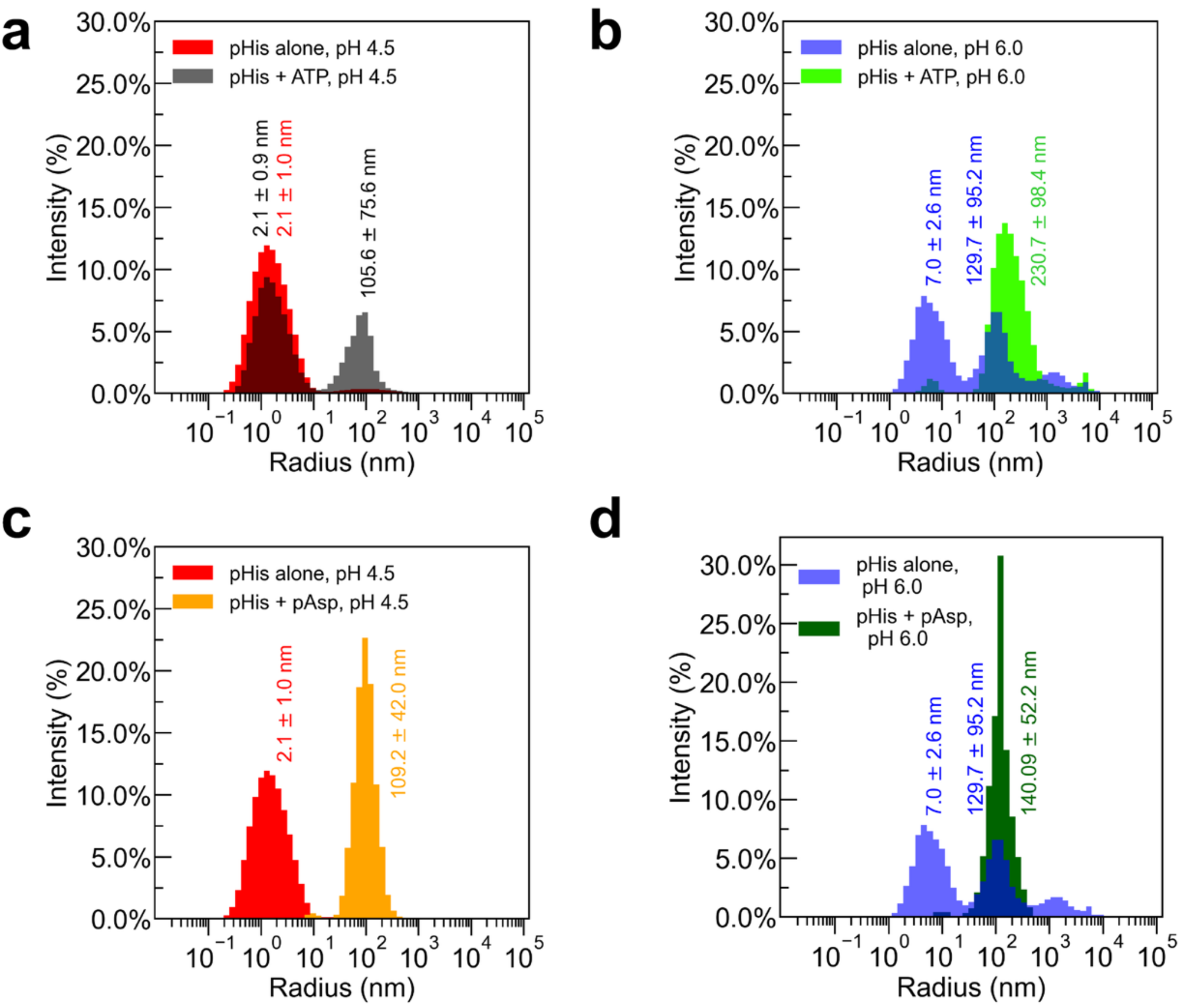

**Supplementary Figure 13.** Dynamic light scattering of pHis complexed with ATP **(a, b)** or pAsp **(c, d)** at either pH 4.5 or pH 6.0. pHis was fixed at 10 mM monomer for all trials. ATP was fixed at 33 μM final concentration and pAsp was fixed at 100 μM monomer as applicable. Attempts to collect ATP and pAsp only measurements at these concentrations yielded poor data quality, presumably because of the low concentration. Intensity values were used in analysis to mitigate assumptions necessary for mass % conversion.

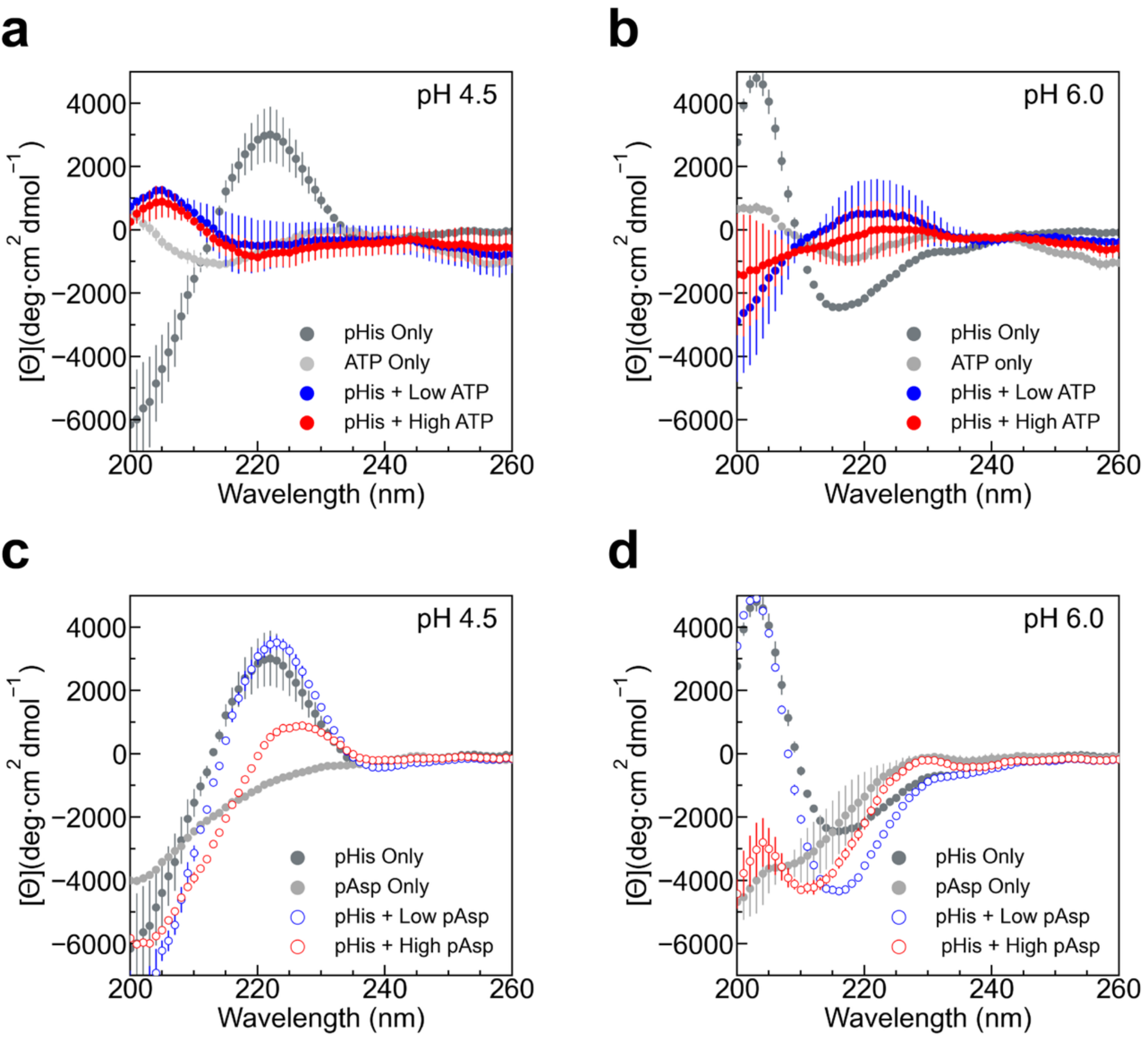

**Supplementary Figure 14**. Circular dichroism spectra of pHis alone and with ATP **(a, b)** or pAsp **(c, d)** at varying pH. CD spectra normalized to mean residue ellipticity ([Θ]) as described in methods. pHis was fixed at 2 mM monomer concentration. Samples containing ATP (a and b) were prepared at low (0.125 mM, blue curve) or high (0.5 mM ATP, red curve) [ATP]. Samples containing pAsp **(c, d)** were prepared at low (0.5 mM monomer, blue curve) or high (2 mM monomer, red curve) [pAsp]. Control spectra of ATP and pAsp alone at either pH were collected at their respective high concentrations and plotted corresponding to those conditions for comparison (light grey respectively).

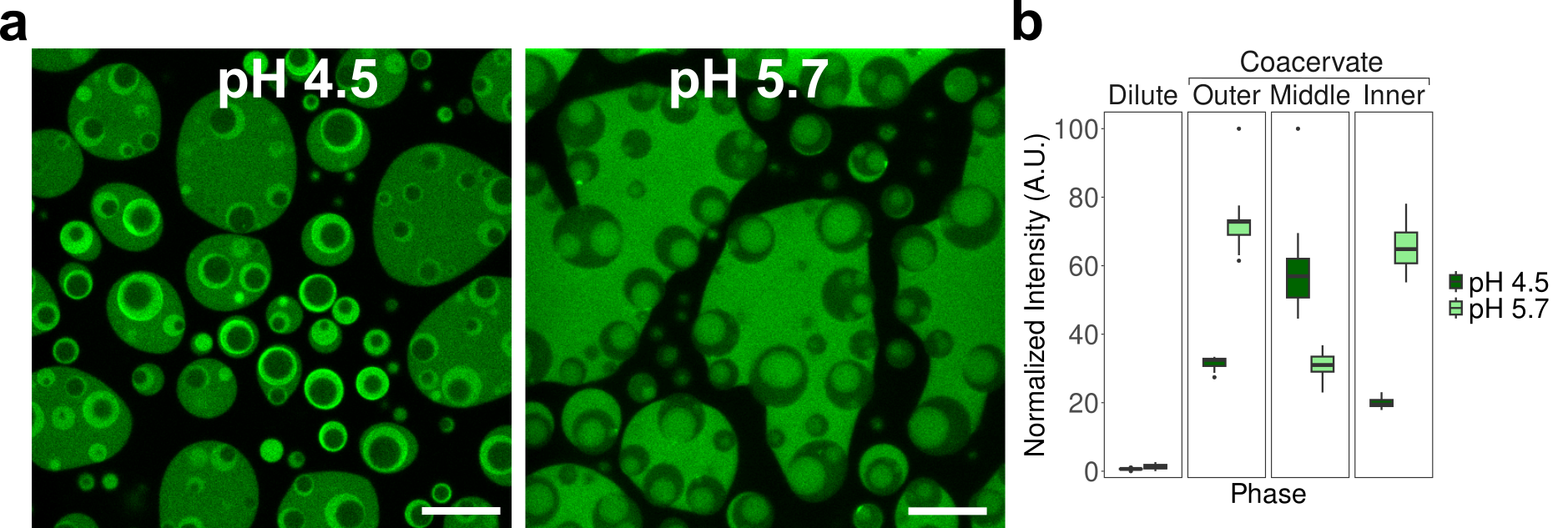

**Supplementary Figure 15.** Distribution of labeled pAsp in the multiphase coacervate system. **(a)** Confocal microscopy images of DEAE-dextran/CM-dextran, pLys/pAsp, and pHis/pAsp coacervates at pH 4.5 and 5.7. pAsp was labeled with Alexa Fluor 488 (green). **(b)** Box plots show the normalized fluorescence intensity (min–max scaling) of pAsp in each phase at both pH values; each dataset was normalized independently. A minimum of 20 intensity measurements were collected per phase. All scale bars: 10 µm.

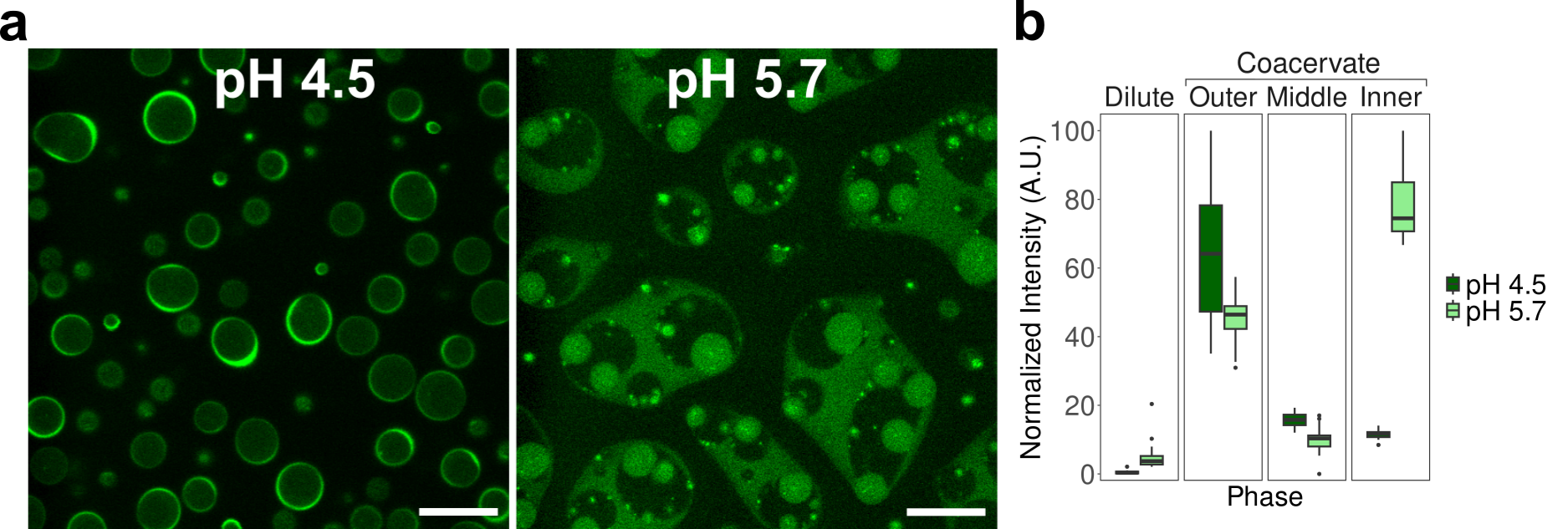

**Supplementary Figure 16.** Distribution of labeled CM-dextran in the multiphase coacervate system. **(a)** Confocal microscopy images of DEAE-dextran/CM-dextran, pLys/pAsp, and pHis/pAsp coacervates at pH 4.5 and 5.7. CM-dextran was labeled with Alexa Fluor 488 (green). **(b)** Box plots show the normalized fluorescence intensity (min–max scaling) of CM-dextran in each phase at both pH values; each dataset was normalized independently. A minimum of 20 intensity measurements were collected per phase. All scale bars: 10 µm.

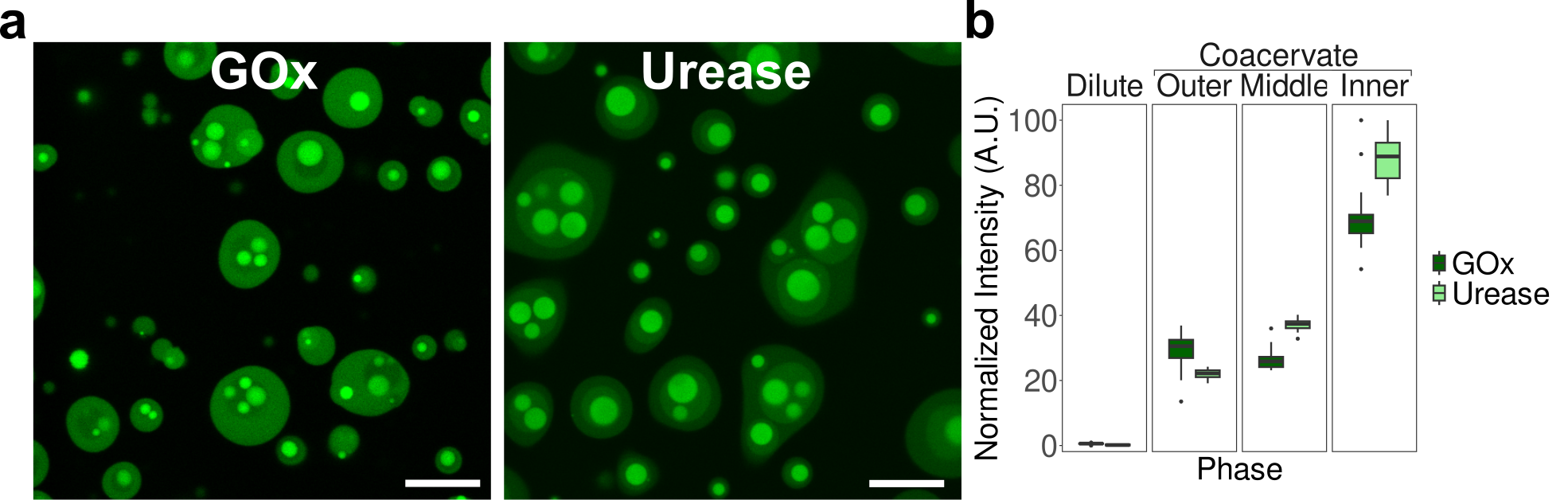

**Supplementary Figure 17.** Distribution of labeled enzymes in a multiphase coacervate system. **(a)** Confocal microscopy images of DEAE-dextran/CM-dextran, pLys/pAsp, and pHis/pAsp coacervates at pH 4.5 containing either urease or glucose oxidase (GOx), each labeled with Alexa Fluor 488 (green). **(b)** Box plots show the normalized fluorescence intensity (min–max scaling) of urease or GOx within each phase; each enzyme system was normalized independently. A minimum of 20 intensity measurements were collected per phase. All scale bars: 10 µm.

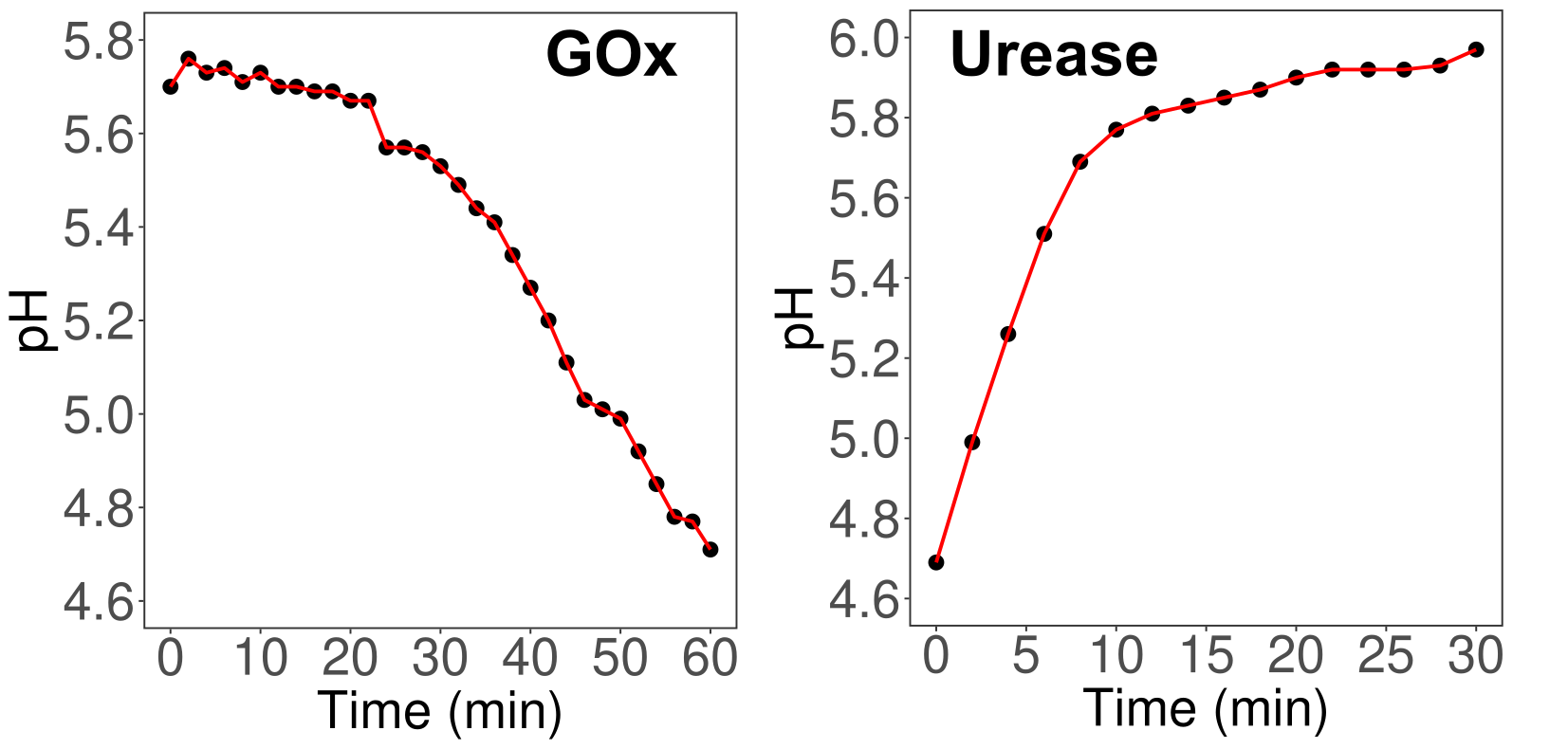

**Supplementary Figure 18.** Enzyme-mediated pH changes over time for the DEAE-dextran/CM-dextran, pLys/pAsp, and pHis/pAsp multiphase coacervate system containing either 0.2 mg/mL glucose oxidase (GOx) or 0.2 mg/mL urease. The GOx reaction was initiated by the addition of 15 mM glucose, and the urease reaction was initiated by the addition of 5 mM urea. Buffer concentrations were 10 mM MES for the GOx experiment and 10 mM acetate for the urease experiment. Both experiments were performed as bulk measurements, with the pH probe inserted directly into the coacervate suspension. Figure labels indicate the corresponding enzyme used in each condition.

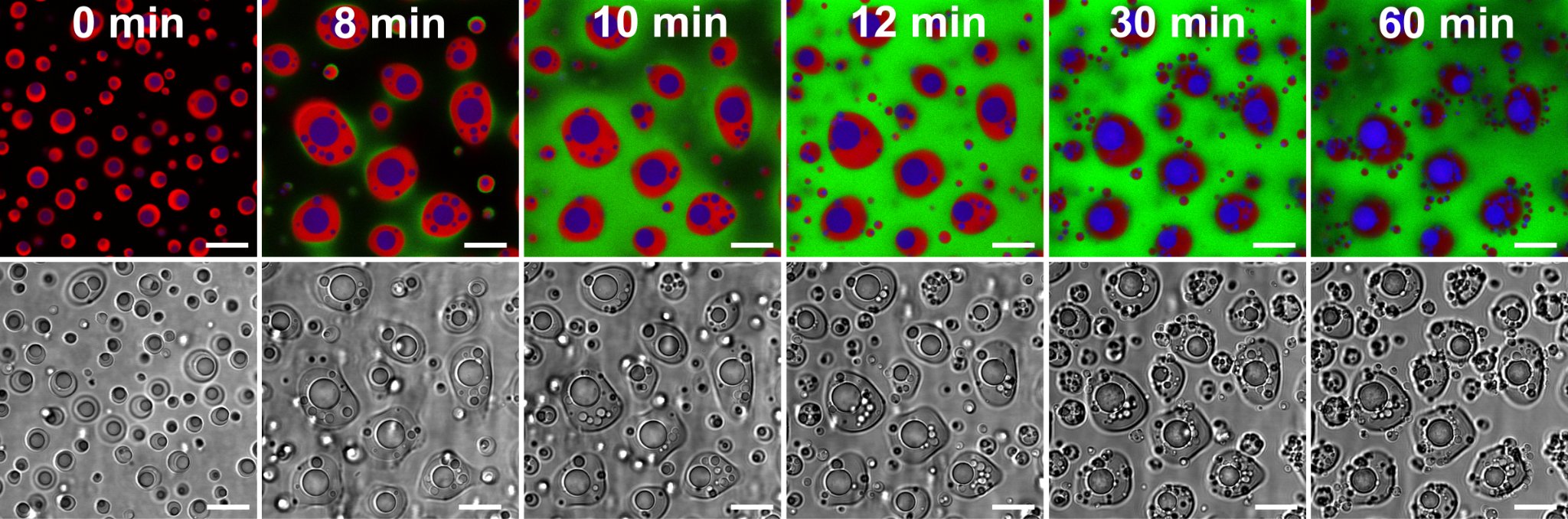

**Supplementary Figure 19.** Phase reorganization driven by urease-mediated pH increase. Confocal microscopy images of DEAE-dextran/CM-dextran, pLys/pAsp, and pHis/pAsp coacervates containing 0.2 mg.mL^-1^ urease, imaged at pH 4.7 before urea addition (0 min, left). The reaction was initiated by the addition of 5 mM urea, resulting in a pH increase to approximately 6.2 after 60 minutes. DEAE-dextran was labeled with fluorescein (green), pLys with rhodamine (red), and pHis with Alexa Fluor 633 (blue). Separate fluorescence channels were false-colored and merged for clarity. Scale bars: 10 µm.

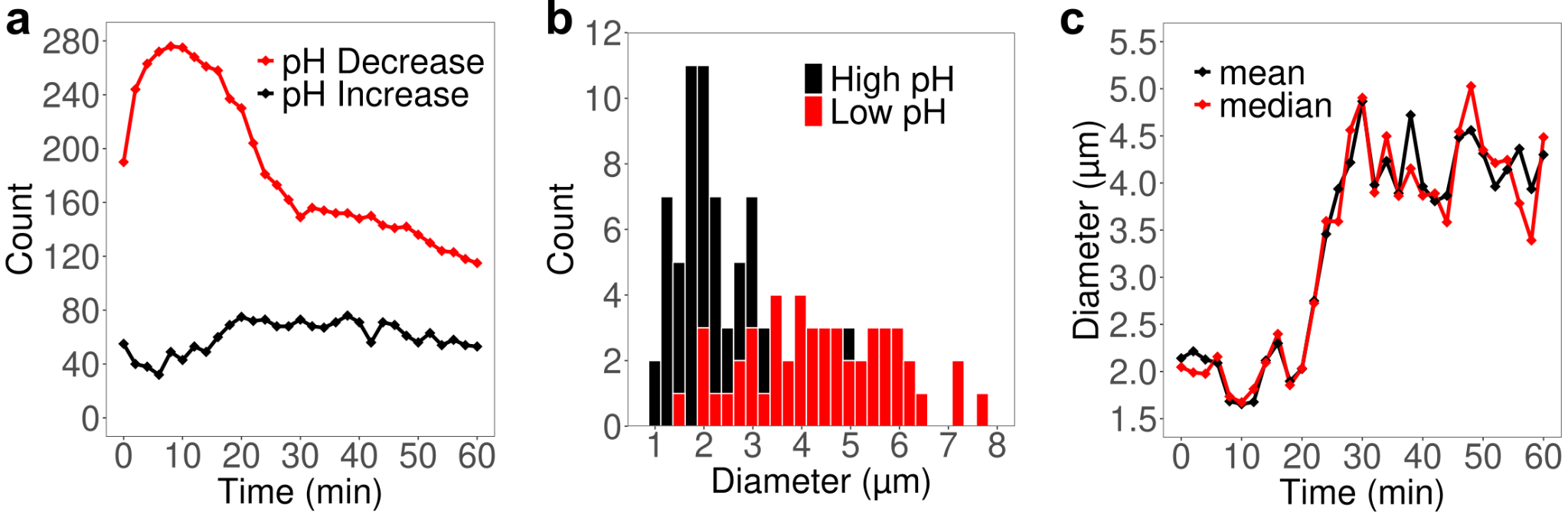

**Supplementary Figure 20.** Plots showing changes in the physical characteristics of the multiphase coacervate system as a function of pH. **(a)** Number of pHis coacervate droplets observed as the pH decreases (GOx and glucose) or increases (urease and urea). **(b)** Histograms of pHis droplet diameters at high (pH 6.2) and low (pH 4.7) conditions. **(c)** Mean and median diameters of pHis droplets as the pH is decreased using GOx and glucose.

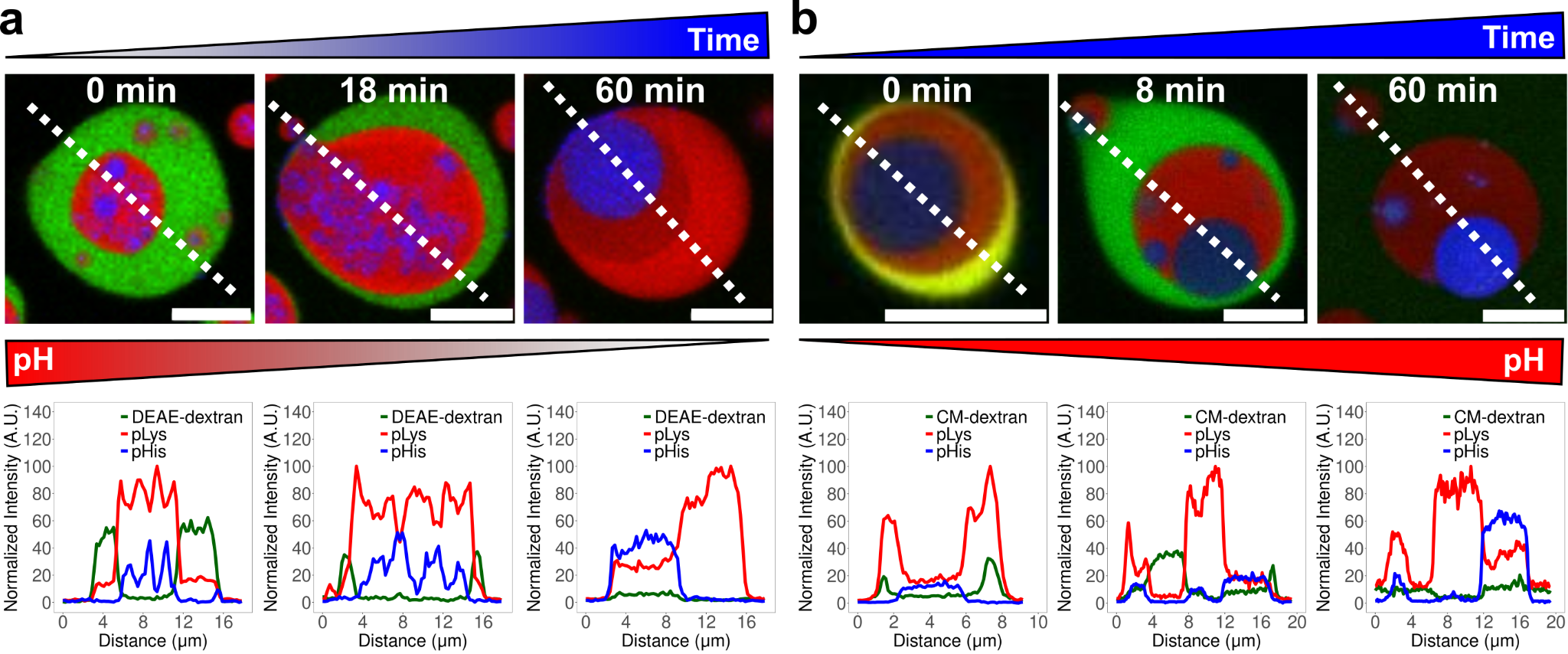

**Supplementary Figure 21. (a, b)** Confocal microscopy images and corresponding line profiles of DEAE-dextran/CM-dextran, pLys/pAsp, and pHis/pAsp coacervates containing 0.2 mg.mL^-1^ of either GOx **(a)** or urease **(b)**. Reactions were initiated by the addition of 15 mM glucose for GOx **(a)** or 5 mM urea for urease (b). In **(a)**, DEAE-dextran was labeled with fluorescein (green), pLys with rhodamine (red), and pHis with Alexa Fluor 633 (blue). In **(b)**, CM-dextran was labeled with Alexa Fluor 488, pLys with rhodamine, and pHis with Alexa Fluor 633. Scale bars: 10 µm.

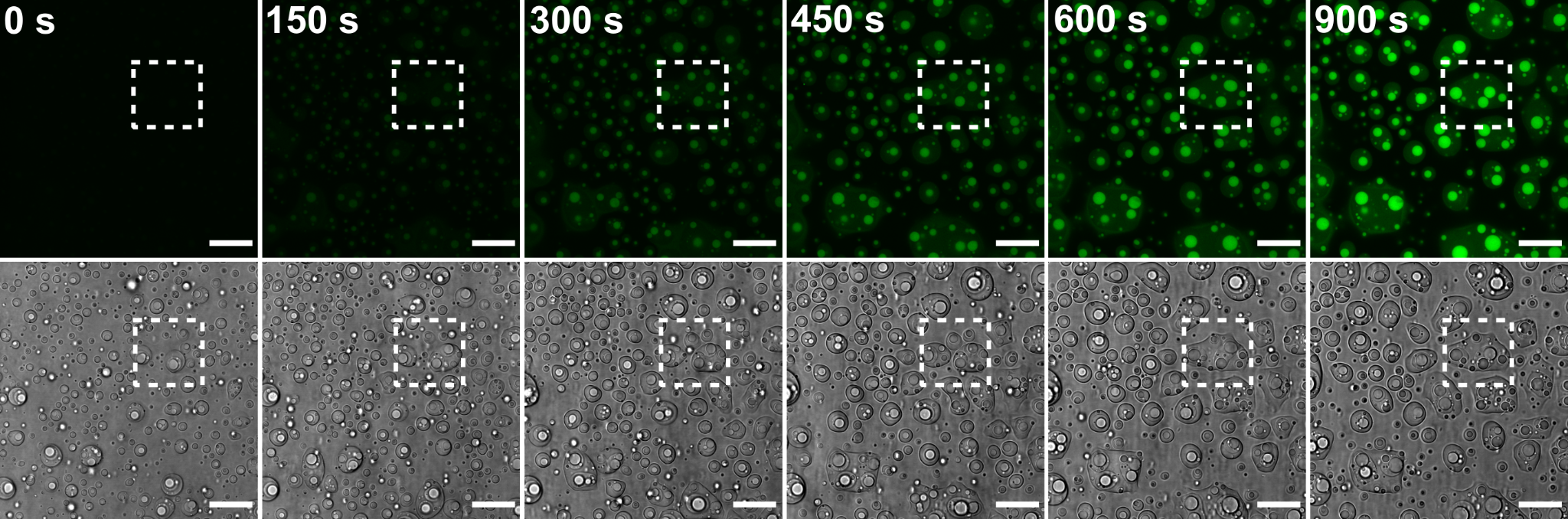

**Supplementary Figure 22.** Confocal microscopy images of DEAE-dextran/CM-dextran, pLys/pAsp and pHis/pAsp coacervates showing 5(6)-Carboxy-2′,7′-dichlorofluorescein diacetate (CDFDA) hydrolysis over time at pH 5.7 to 5(6)-Carboxy-2′,7′-dichlorofluorescein (green). Dashed line box corresponds to region depicted in Fig. 5d in main text. Scale bars = 20 µm.

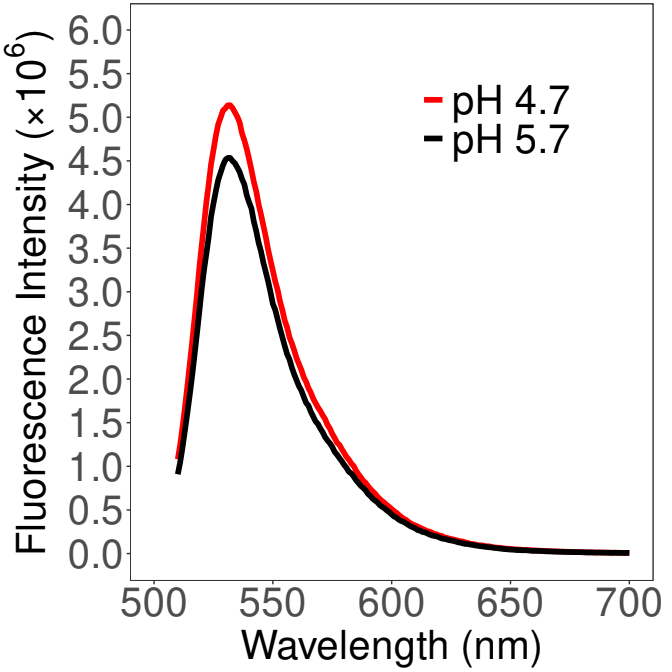

**Supplementary Figure 23.** Fluorescence spectra of 0.4 µM 5(6)-Carboxy-2′,7′-dichlorofluorescein (CDF) in DEAE-dextran/CM-dextran, pLys/pAsp, and pHis/pAsp multiphase coacervates at pH 4.7 and 5.7. The fluorescence intensity at 529 nm was 1.14-fold higher at pH 5.7 than at pH 4.7, and this ratio was used as a correction factor in all subsequent fluorescence spectroscopy kinetics experiments (Fig. 5f–h, Fig. 6d, and Supplementary Figs. 24, 25, and 30 and Supplementary Table 2). This spectrum was also used to convert fluorescence units to concentration values in kinetic fluorescence spectroscopy.

| **Coacervate** | **pH** | **Rate (nMs^-1^)** | **Estimated % Complete** |
| --- | --- | --- | --- |
| pAsp Multiphase | pH 4.7 | 0.46 | 3.5 |
| pAsp Multiphase | pH 5.7 | 3.4 | 21 |
| ATP Multiphase | pH 4.7 | 0.020 | 0.15 |
| ATP Multiphase | pH 5.7 | 0.040 | 1.0 |
| pAsp Single-phase | pH 4.7 | 0.52 | 6.3 |
| pAsp Single-phase | pH 5.7 | 0.080 | 0.92 |
| ATP Single-phase | pH 4.7 | 0.00 | 0.15 |
| ATP Single-phase | pH 5.7 | 0.00 | 1.6 |
| No pHis | pH 4.7 | 0.010 | 0.030 |
| No pHis | pH 5.7 | 0.010 | 0.11 |
| No Polymers | pH 4.7 | 0.010 | 0.15 |
| No Polymers | pH 5.7 | 0.050 | 0.33 |
| pHis Only | pH 4.7 | 9.2 | 40 |
| pHis Only | pH 5.7 | 9.3 | 31 |
| Adenosine | pH 5.7 | 2.6 | 17 |
| AMP | pH 5.7 | 6.6 | 22 |
| ADP | pH 5.7 | 1.9 | 9.6 |
| Urease | N/A | 3.8 | 21 |

### Supplementary Table 2.

**Conditions of the fluorescence spectroscopy kinetics experiments and percentage completion of CDFDA hydrolysis to CDF in the coacervate system.** The relationship between fluorescence intensity and CDF concentration (Supplementary Fig. 23) was used to convert fluorescence signals to concentration, enabling calculation of hydrolysis rates over the first 30 s. Percent completion was determined by comparing the final measured fluorescence intensity to the theoretical maximum, assuming complete conversion of CDFDA to CDF.

**Table definitions:**

- **pAsp multiphase**: DEAE-dextran/CM-dextran, pLys/pAsp, and pHis/pAsp multiphase system.
- **ATP multiphase**: Same as pAsp multiphase, except ATP in pHis/pAsp is replaced with ATP.
- **pAsp single-phase** / **ATP single-phase**: pHis/pAsp or pHis/ATP single-phase systems, respectively.
- **No pHis**: CM-dextran/DEAE-dextran and pLys/pAsp multiphase coacervate without pHis.
- **No polymers**: Buffer only, no coacervates.
- **pHis only**: pHis in solution without partner ions or coacervates.
- **Adenosine**, **AMP**, **ADP**: ATP in a CM-dextran/DEAE-dextran, pLys/pAsp, and pHis/ATP multiphase system replaced with an equal concentration of the indicated compound.
- **Urease**: Urease reaction (5 mM urea) in a DEAE-dextran/CM-dextran, pLys/pAsp, and pHis/pAsp multiphase system.

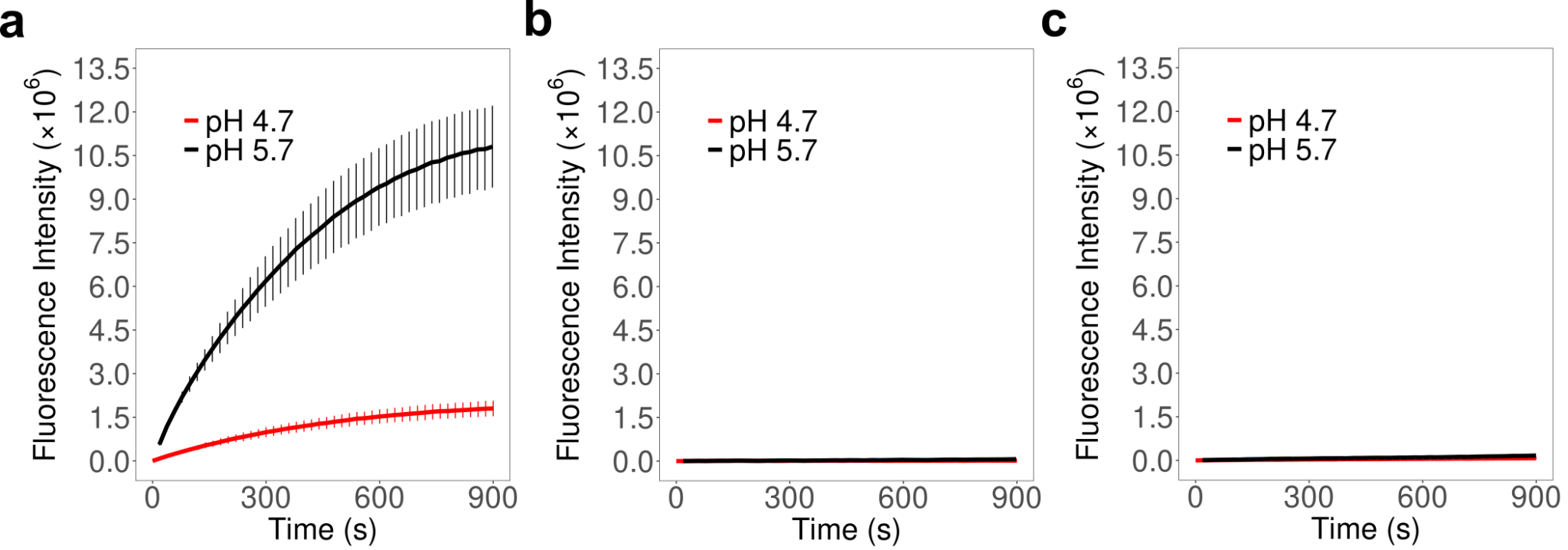

**Supplementary Figure 24.** **(a–c)** Fluorescence spectroscopy kinetics showing the hydrolysis of 5 µM CDFDA in: **(a)** DEAE-dextran/CM-dextran, pLys/pAsp, and pHis/pAsp multiphase coacervates (without ATP); **(b)** DEAE-dextran/CM-dextran and pLys/pAsp multiphase coacervates (without pHis); and **(c)** buffer only (no coacervates).

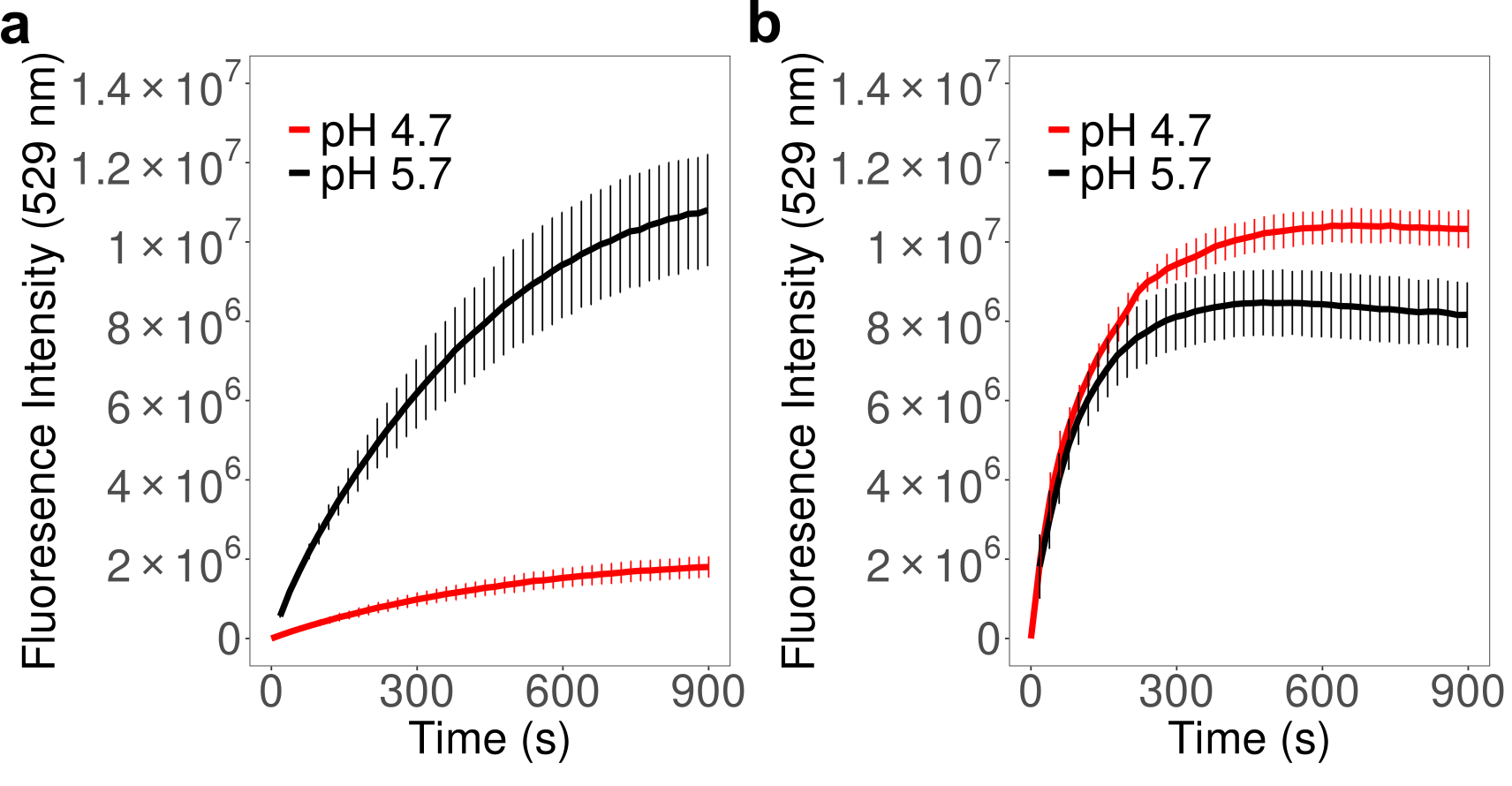

**Supplementary Figure 25.** **(a, b)** Fluorescence spectroscopy kinetics showing the hydrolysis of CDFDA at pH 4.7 and 5.7 in: **(a)** DEAE-dextran/CM-dextran, pLys/pAsp, and pHis/pAsp multiphase coacervates (without ATP); and **(b)** pHis (10 µM) alone, without coacervates. To keep fluorescence within the detector’s linear range, experiments in **(a)** were performed with 5 µM CDFDA, and those in **(b)** with 2.5 µM CDFDA.

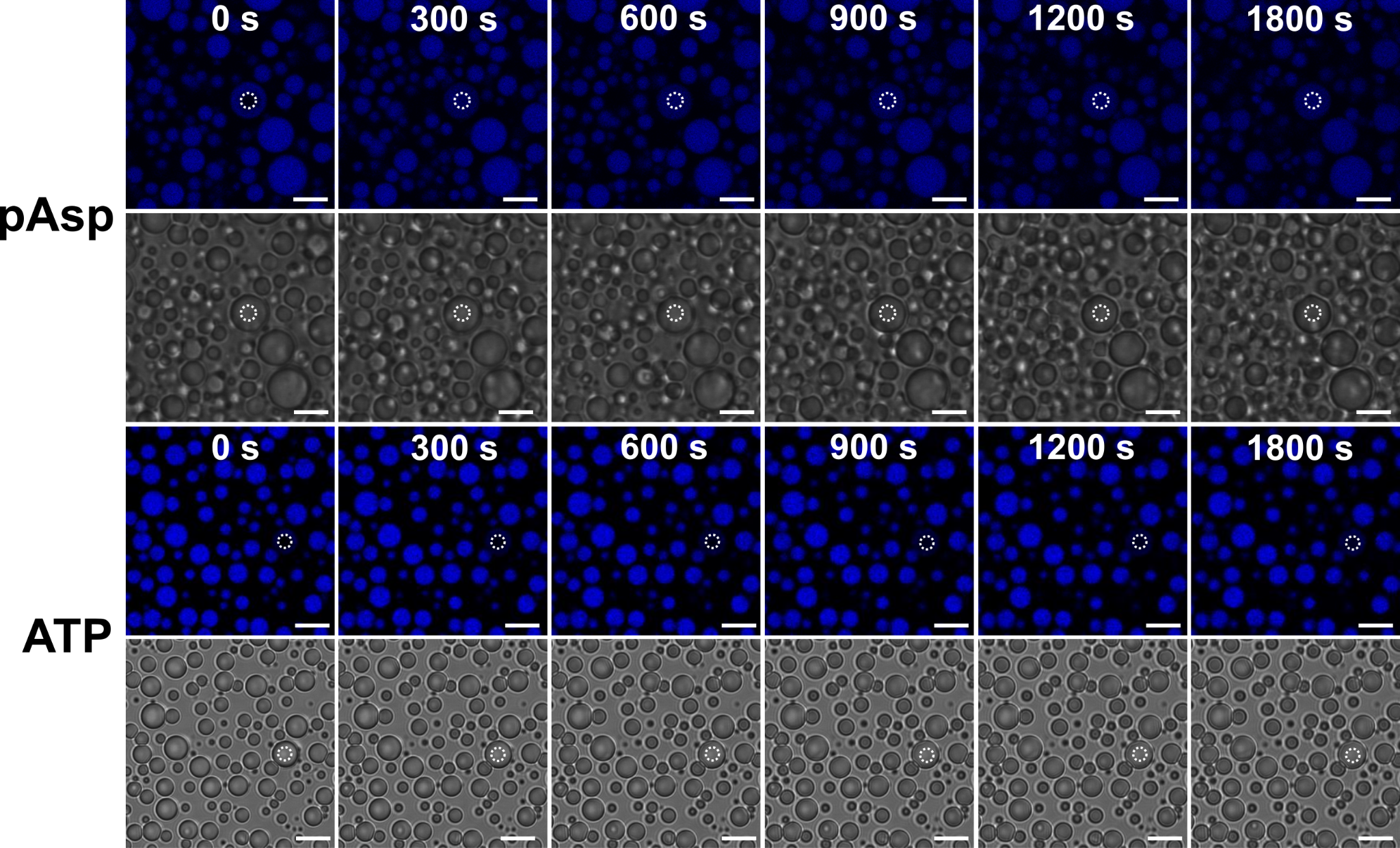

**Supplementary Figure 26.** Confocal microscopy images showing the fluorescence recovery after photobleaching of pHis/pAsp or pHis/ATP single-phase coacervates at pH 5.7 (pHis is labelled with Alexa Fluor 633, which is blue). All images are false-colored for clarity. All scale bars = 10 µm.

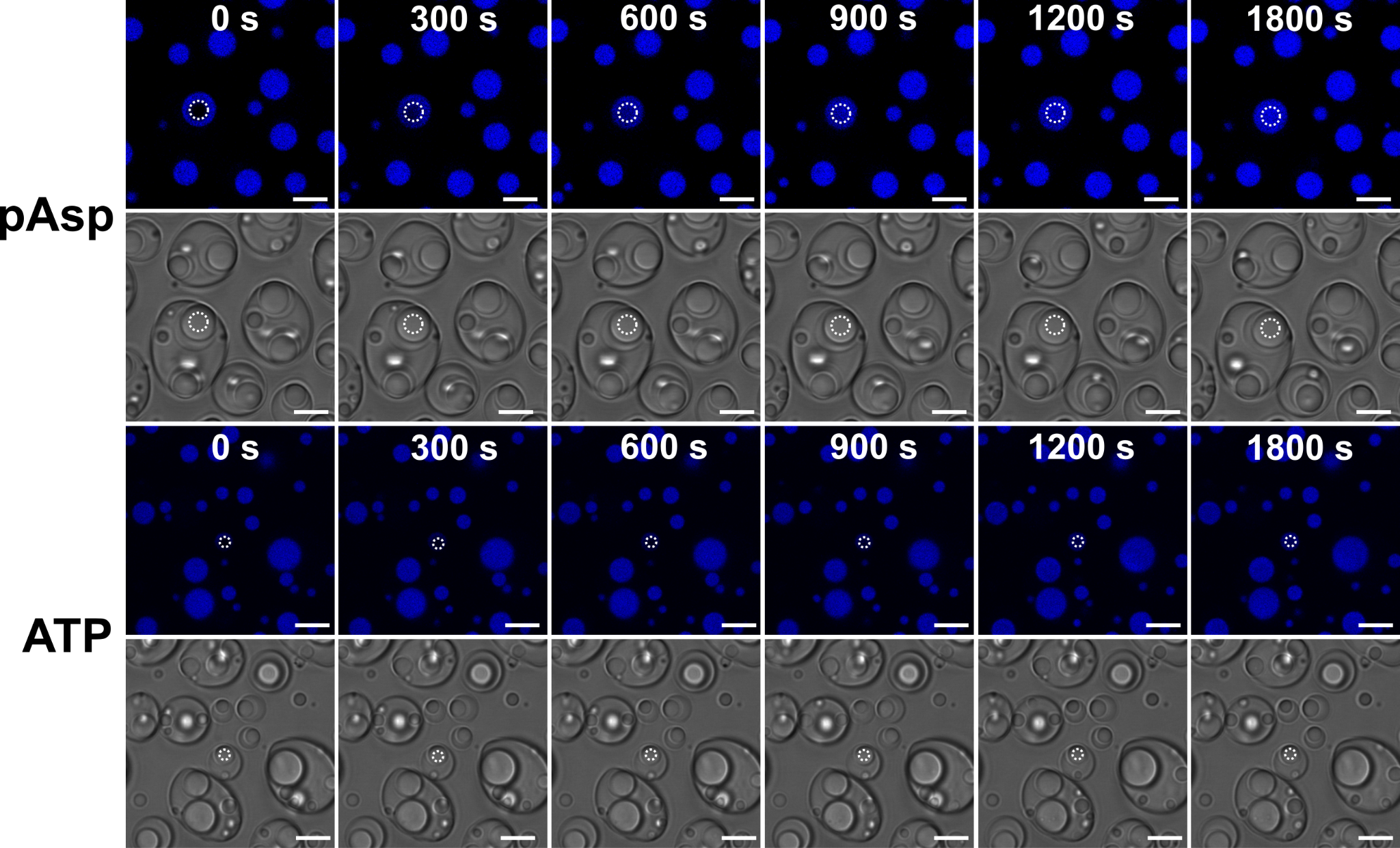

**Supplementary Figure 27.** Confocal microscopy images showing the fluorescence recovery after photobleaching of pHis rich coacervate phase (labeled with Alexa Fluor 633) contained within a multiphase coacervate system. Top two rows consist of DEAE-dextran/CM-dextran, pLys/pAsp and pHis/pAsp coacervates at pH 5.7. Bottom two rows consist of DEAE-dextran/CM-dextran, pLys/pAsp and pHis/ATP coacervates at pH 5.7. In both systems, pHis was labelled with Alexa Fluor 633 (blue) and images are false-colored for clarity. All scale bars = 10 µm.

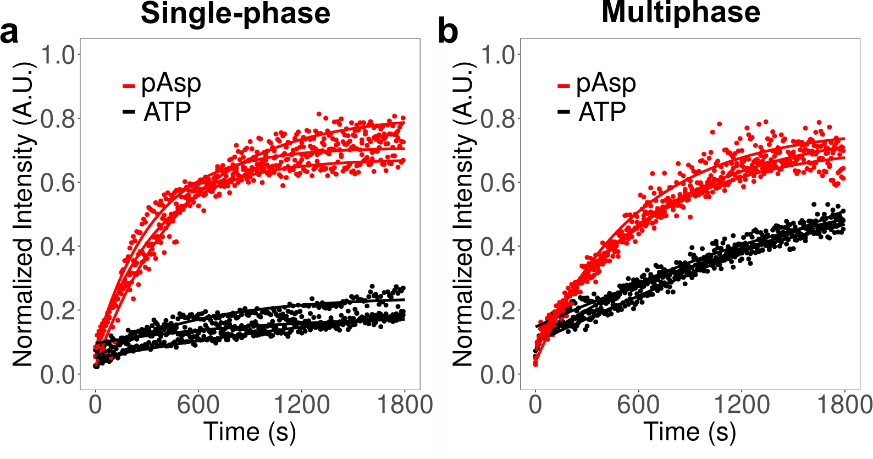

**Supplementary Figure 28. (a, b)** Fluorescence recovery after photobleaching (FRAP) of the pHis-rich phase (labelled with Alexa Fluor 633) in **(a)** pHis/pAsp and pHis/ATP single-phase systems, and **(b)** DEAE-dextran/CM-dextran, pLys/pAsp, and either pHis/pAsp or pHis/ATP multiphase systems, all at pH 5.7. In both cases, the pAsp-complexed systems showed faster recovery, with the difference being more pronounced in the single-phase system than in the multiphase system. Panel **(a)** corresponds to microscopy images in Supplementary Fig. 26, and panel **(b)** to those in Supplementary Fig. 27.

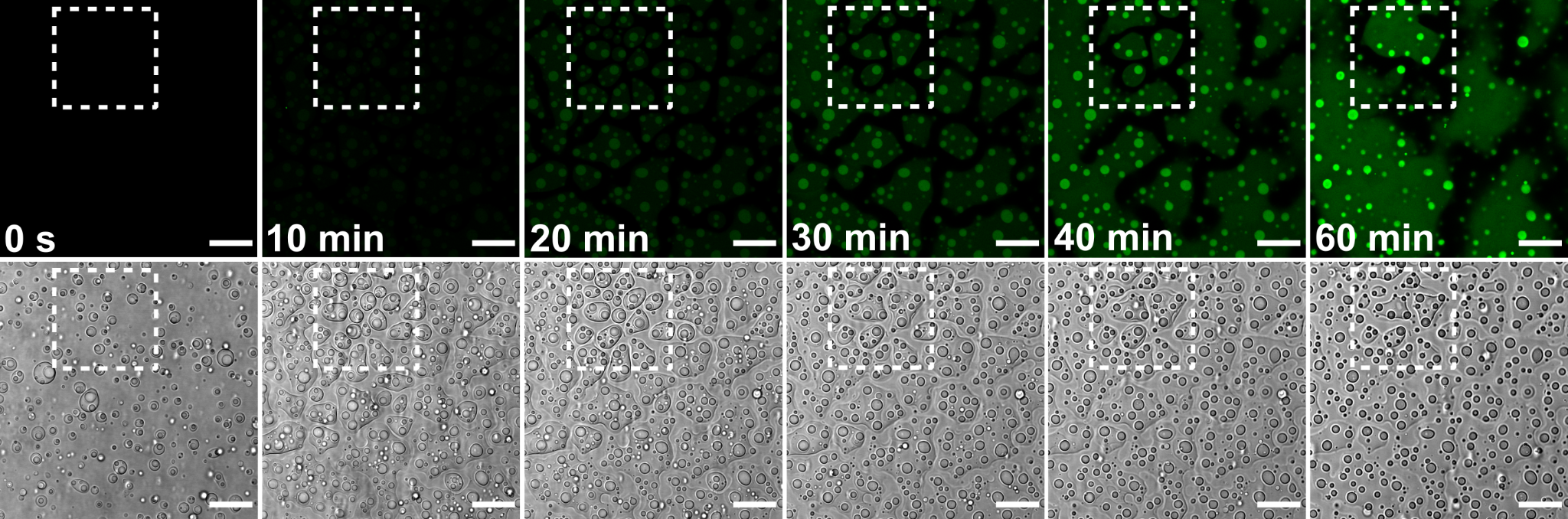

**Supplementary Figure 29.** Confocal microscopy images showing CDFDA hydrolysis in DEAE-dextran/CM-dextran, pLys/pAsp, and pHis/pAsp coacervates. The system was initially at pH 4.7, which increased to pH 6.5 over the course of the experiment through urease-mediated hydrolysis of urea. The dashed box indicates the region shown in Fig. 6b of the main text. Scale bars: 20 µm.

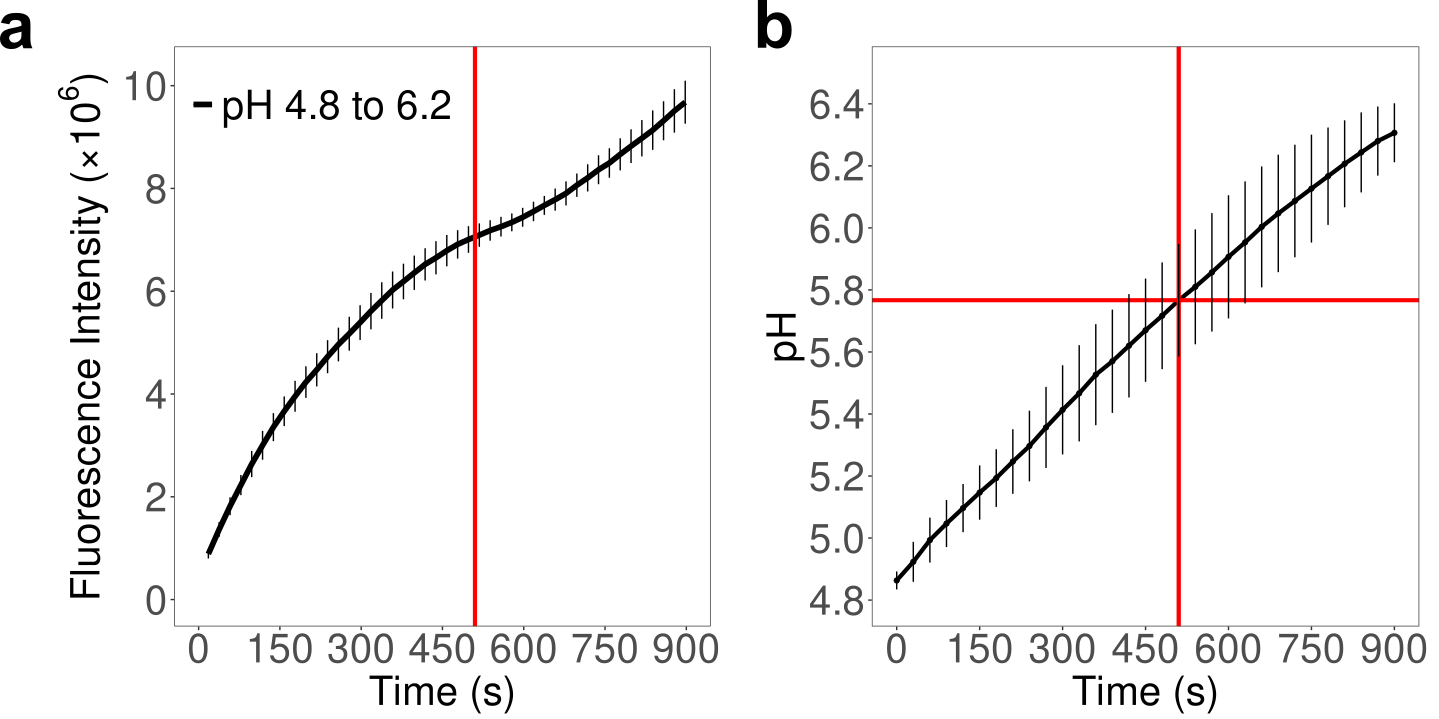

**Supplementary Figure 30. (a)** Fluorescence spectroscopy kinetics showing the hydrolysis of CDFDA in the DEAE-dextran/CM-dextran, pLys/pAsp, and pHis/pAsp multiphase system, in which the pH increases from 4.8 to 6.2 over 900 seconds due to urease-mediated hydrolysis of 5 mM urea (λ = 529 nm). The corresponding pH change (**b**) was measured by inserting a pH electrode directly into the multiphase coacervate suspension. The red line indicates the time point (510 s) at which the rate of fluorescence intensity change begins to level off in the S-shaped plot (**a**); this corresponds to a measured pH of 5.76 (**b**).
